# Consistency across multi-omics layers in a drug-perturbed gut microbial community

**DOI:** 10.1101/2023.01.03.519475

**Authors:** Sander Wuyts, Renato Alves, Maria Zimmermann-Kogadeeva, Suguru Nishijima, Sonja Blasche, Marja Driessen, Philipp E. Geyer, Rajna Hercog, Ece Kartal, Lisa Maier, Johannes B. Müller, Sarela Garcia Santamarina, Thomas Sebastian B. Schmidt, Daniel C. Sevin, Anja Telzerow, Peter V. Treit, Tobias Wenzel, Athanasios Typas, Kiran R. Patil, Matthias Mann, Michael Kuhn, Peer Bork

**Affiliations:** European Molecular Biology Laboratory, Heidelberg, Germany; Medical Research Council Toxicology Unit, Cambridge, UK; Department of Proteomics and Signal Transduction, Max Planck Institute of Biochemistry, Martinsried, Germany; Cellzome, GlaxoSmithKline R&D, Heidelberg, Germany; Proteomics Program, NNF Center for Protein Research, Faculty of Health Sciences, University of Copenhagen, Copenhagen, Denmark; Max Delbrück Centre for Molecular Medicine, Berlin, Germany; Yonsei Frontier Lab (YFL), Yonsei University, Seoul, South Korea; Department of Bioinformatics, Biocenter, University of Würzburg, Würzburg, Germany

## Abstract

Multi-omics analyses are increasingly employed in microbiome studies to obtain a holistic view of molecular changes occurring within microbial communities exposed to different conditions. However, it is not always clear to what extent each omics data type contributes to our understanding of the community dynamics and whether they are concordant with each other. Here we map the molecular response of a synthetic community of 32 human gut bacteria to three non-antibiotic drugs by using five omics layers, namely 16S rRNA gene profiling, metagenomics, metatranscriptomics, metaproteomics, and metabolomics. Using this controlled setting, we find that all omics methods with species resolution in their readouts are highly consistent in estimating relative species abundances across conditions. Furthermore, different omics methods complement each other in their ability to capture functional changes in response to the drug perturbations. For example, while nearly all omics data types captured that the antipsychotic drug chlorpromazine selectively inhibits Bacteroidota representatives in the community, the metatranscriptome and metaproteome suggested that the drug induces stress responses related to protein quality control and metabolomics revealed a decrease in polysaccharide uptake, likely caused by Bacteroidota depletion. Taken together, our study provides insights into how multi-omics datasets can be utilised to reveal complex molecular responses to external perturbations in microbial communities.

## Introduction

The human gut microbiota is a complex community of microorganisms, which is affected by endogenous and environmental factors such as host genotype, diet, drug treatment, and disease status, and in turn influences host health and disease progression (Kau *et al,* 2011; Cho & Blaser, 2012; Cani, 2018; Durack & Lynch, 2018; Schmidt *et al,* 2018; Lindell *et al,* 2022). Currently, insights into structure and function of the microbiota community mainly come from 16S rRNA gene profiling and shotgun metagenomics. While 16S rRNA amplicon sequencing offers a cost-efficient way to assess bacterial abundance at a higher taxonomic level, whole-genome shotgun metagenomics resolves abundance of species and strains, together with the functional potential they encode (Quince *et al,* 2017; Almeida *et al,* 2019; Pasolli *et al,* 2019). In addition, gene and protein expression and metabolite abundance in the community can be quantified with metatranscriptomics (Bashiardes *et al*, 2016), metaproteomics (Zhang & Figeys, 2019) and metabolomics (Zierer *et al,* 2018; Han *et al,* 2021), respectively. Ultimately, the combination of these methods should enable integration of the major molecular layers of the cell, resulting in a more complete picture of the microbiome (Jansson & Baker, 2016; Heintz-Buschart & Wilmes, 2018). Several studies have shown how a combination of two or more of these omics methods could lead to novel insights regarding the dynamics and inner workings of a microbial community (Heintz-Buschart *et al,* 2016; Lloyd-Price *et al*, 2017; Salazar *et al*, 2019; Taylor *et al*, 2020). While multi-omics measurements provide information across molecular layers, their comprehensive integration remains challenging. One challenge is the limited knowledge about the concordance of different measurements in complex *in natura* settings in the absence of ground truth. Another challenge in comparing and integrating multi-omics datasets is the difference in their dynamics in response to perturbations. Whereas metabolite changes occur on a time scale of seconds, transcriptional changes usually occur on a time scale of minutes, while protein abundance changes take the longest to respond to a perturbation (Gerosa & Sauer, 2011; Choi *et al,* 2020).

Synthetic microbial communities have been increasingly used to obtain a better understanding of the dynamics and species–species interactions (Goldford *et al,* 2018; Cheng *et al,* 2021). Compared to a natural gut microbiota, these synthetic communities have lower complexity, higher controllability and reproducibility, and a well-defined composition at the strain level, at the cost of being simplified representations of natural ecosystems (Roy *et al,* 2014; Weiss *et al,* 2022; Aranda-Díaz *et al,* 2022). Yet, they do offer advantages over single species studies, as single species’ behaviour can significantly differ in mono-culture compared to co-culture (D’hoe *et al,* 2018).

The complex interactions between the gut microbiota and non-antibiotic drugs have been elucidated from large-scale human studies and high-throughput laboratory experiments (Rizkallah & Aziz, 2010; Forslund *et al,* 2015; Spanogiannopoulos *et al,* 2016; Wilson & Nicholson, 2017; Zimmermann *et al,* 2021; Forslund *et al,* 2021). This relationship is bidirectional, as drugs can influence microbiome composition (Maier *et al,* 2018; Jackson *et al,* 2018; Vich Vila *et al,* 2020; Vieira-Silva *et al,* 2020), while the gut microbiota can have an impact on a drug’s efficacy and toxicity by altering its chemical structure (Zimmermann *et al,* 2019a, 2019b; Javdan *et al,* 2020; Klünemann *et al,* 2021). The emerging knowledge on drug–microbiota interactions has the potential to influence the future of drug development and personalized medicine (Doestzada *et al,* 2018; Weersma *et al,* 2020; Maier *et al,* 2021; Zimmermann *et al*, 2021).

To systematically assess and compare how multi-omics measurements capture dynamic changes in microbial communities in response to perturbations, we designed a controlled time-course experiment with a synthetic community of 32 human gut representatives (Tramontano *et al,* 2018) in response to three drugs from diverse indication areas: chlorpromazine (antipsychotic), metformin (antidiabetic) and niclosamide (anthelmintic), which were previously reported to impair growth of several gut bacteria (Maier *et al,* 2018). We followed the response of the defined community to the three non-antibiotic drugs over four days on the structural and functional levels across multi-omics layers, based on 16S rRNA gene, metagenome, metatranscriptome, metaproteome and untargeted metabolome profiling.

## Results

### Establishment of a synthetic community for drug perturbations

To investigate microbial community response to drug perturbations in a controlled system across five omics layers, we combined 32 human gut microbiome representatives (Tramontano *et al*, 2018) and exposed this community to three different non-antibiotic drugs (Figure 1A). The complete experiment was performed twice (run A and run B) as biological replicates, starting from the initial community assembly step from single bacterial cultures. More specifically, seven slow-growing species (inoculated on day 1) were combined with 25 fast-growing species (inoculated on day 3) on day 5 to form a synthetic community (Figure 1A, B). In order to ensure stable community composition, we performed three culture passages by growing the mixed culture for 48 hours and transferring 1% of total volume to a fresh culture medium. Samples for 16S rRNA amplicon sequencing were taken immediately after combining the strains (Inoculum mix) and after each passage (Transfer 1 - 3) to evaluate the stabilisation of the community (Figure 1A; top row). We found that in both runs of the experiment the community reached a stable composition with four highly abundant species after three transfers (relative abundance >10% for *Escherichia coli, Clostridium perfringens, Veillonella parvula* and *Bacteroides thetaiotaomicron,* Supplementary Figure 1A). Bray-Curtis dissimilarity index showed that both runs were highly similar after the third transfer (Supplementary Figure 1B, C).

**Figure 1.**
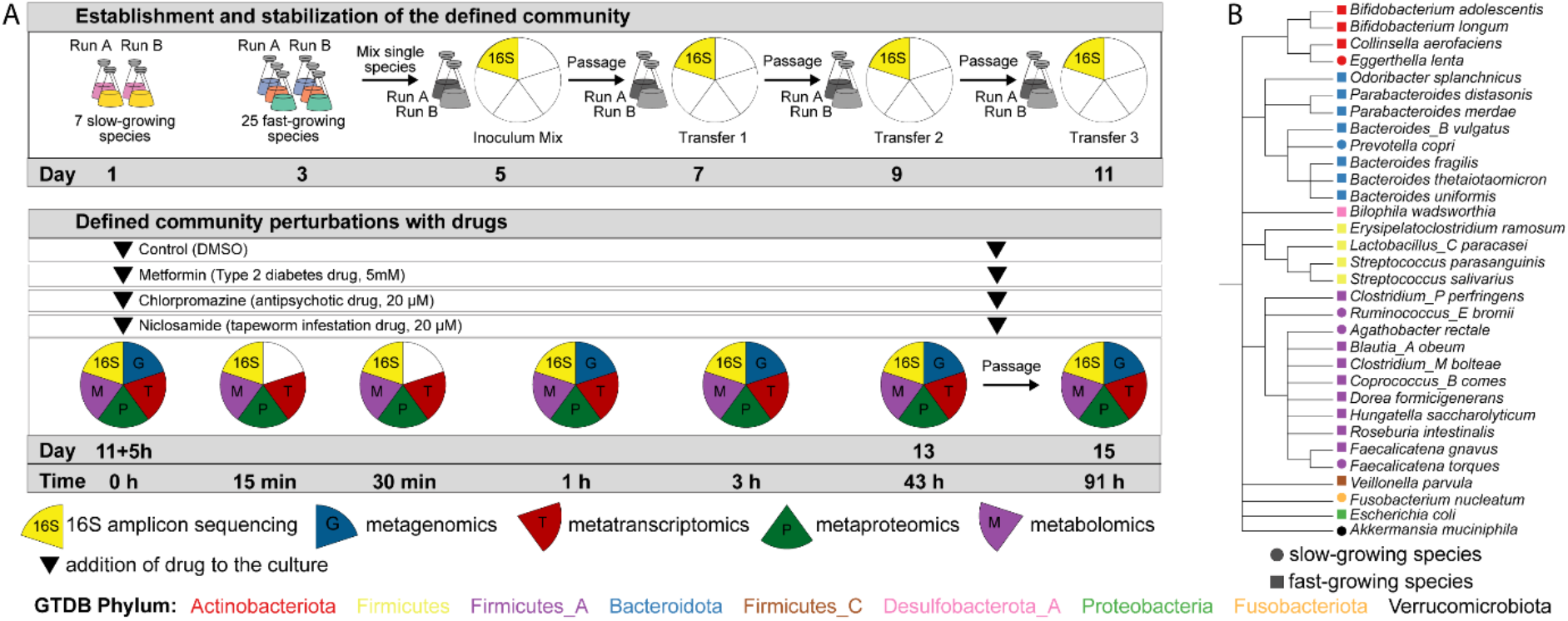
Experimental design and species used in this study. A) Schematic overview of the experimental design. B) Species cladogram constructed by pruning the relevant species from the GTDB species cladogram (release 95).

After stabilisation, in each run the community perturbation was performed in duplicate during exponential growth (i.e., five hours after passaging, as determined by optical density measurements on the previous transfer; Supplementary Figure 1D) by addition of one of the following drugs: i) 5 mM metformin, a type 2 diabetes drug, ii) 20 μM chlorpromazine, an antipsychotic drug, or iii) 20 μM niclosamide, an anthelmintic drug (Figure 1A), while DMSO was used as a control. The higher concentration for metformin was based on reported intestinal concentrations, and previous data on metformin amounts sufficient to impair growth of gut microbiota members *in vitro* (Maier *et al*, 2018; Bailey *et al*, 2008b). The communities were sampled right before the addition of the drugs and 15 min, 30 min, 1 h, and 3 h following the drug perturbation (Figure 1A, Supplementary Table 1). These time points were chosen to elucidate the early response of the bacterial community to drug treatment. After 43 h, an additional sample was taken, and the communities were transferred into a fresh culture medium containing the drugs at initial concentrations. A final sample was taken 48 h after this passage (91 hours after the initial drug addition). In general, high correlation was evident between technical replicates within the same omics dataset (Supplementary Figure 2).

### Consistency of community composition across omics measurements

We first evaluated similarities and differences between the omics measurements in their ability to estimate species abundance. For sequencing-based omics methods, we performed both naïve analyses with commonly used computational pipelines that do not use the information about synthetic community composition (DADA2 for 16S rRNA amplicon sequencing (Callahan *et al,* 2016), mOTUS v2.5 for metagenomics and metatranscriptomics (Milanese *et al,* 2019)), and targeted analyses based on mapping to the 32 reference genomes of species comprising our community (Materials and methods). Within each omics method, both computational approaches produced highly similar results (Supplementary Figure 3). As the composition-naïve approach only yields genus-level resolution for 16S rRNA sequencing data (Knight *et al,* 2018), we used the reference genome mapping approach that yields higher resolution for all methods for comparison of community composition across omics types. For consistency, the same methodology (reference genome mapping) was used for metagenomics and metatranscriptomics. For metaproteomics data, we estimated species abundance by summing protein intensities for all proteins assigned to each species and dividing these values by the total protein intensity in each sample, as suggested previously (Kleiner *et al*, 2017).

We compared relative species abundances between all pairs of omics methods except for metabolomics, which by nature represents total metabolite measurements in the community and does not allow to separate compounds by species. Based on correlation analysis, we found the abundance estimates to be highly similar (minimum Spearman correlation coefficient ρ = 0.78). Congruence was more pronounced for highly abundant species (Figure 2A). Specifically, metagenomics and metatranscriptomics were the most similar of all pairwise comparisons (ρ = 0.92). Further, 16S rRNA amplicon sequencing showed high similarity with metagenomics for species with relative abundances higher than 0.001% (ρ = 0.89). However, for several species with low relative abundances, 16S rRNA sequencing provided higher relative abundance estimates compared to metagenomics, while other species, detected by metagenomics, were not detected with 16S rRNA sequencing. For this observation, no clear taxon-specific or condition-specific effect was found (Supplementary Figure 4), indicating that the differences at these low relative abundances are most likely a result of differences in sequencing depth per sample, as has been previously reported (Pereira-Marques *et al,* 2019; Durazzi *et al,* 2021). Although metaproteomics is not yet widely used for species abundance estimation, we found the corresponding estimates in good agreement with the other omics methods, but only for species with relative abundance above 1% (ρ = 0.78 – 0.84; 16 out of 29 species detected across all samples). This indicates that metaproteomics is less sensitive than sequencing-based methodologies for species abundance estimation, as has also been observed for *in natura* metaproteomics studies (Zhang & Figeys, 2019). Our results show generally high consistency between omics data types in relative species abundance estimations, and underline that metaproteomics can, in principle, provide robust species abundance estimates, at least for synthetic microbial communities, albeit with lower sensitivity.

**Figure 2.**
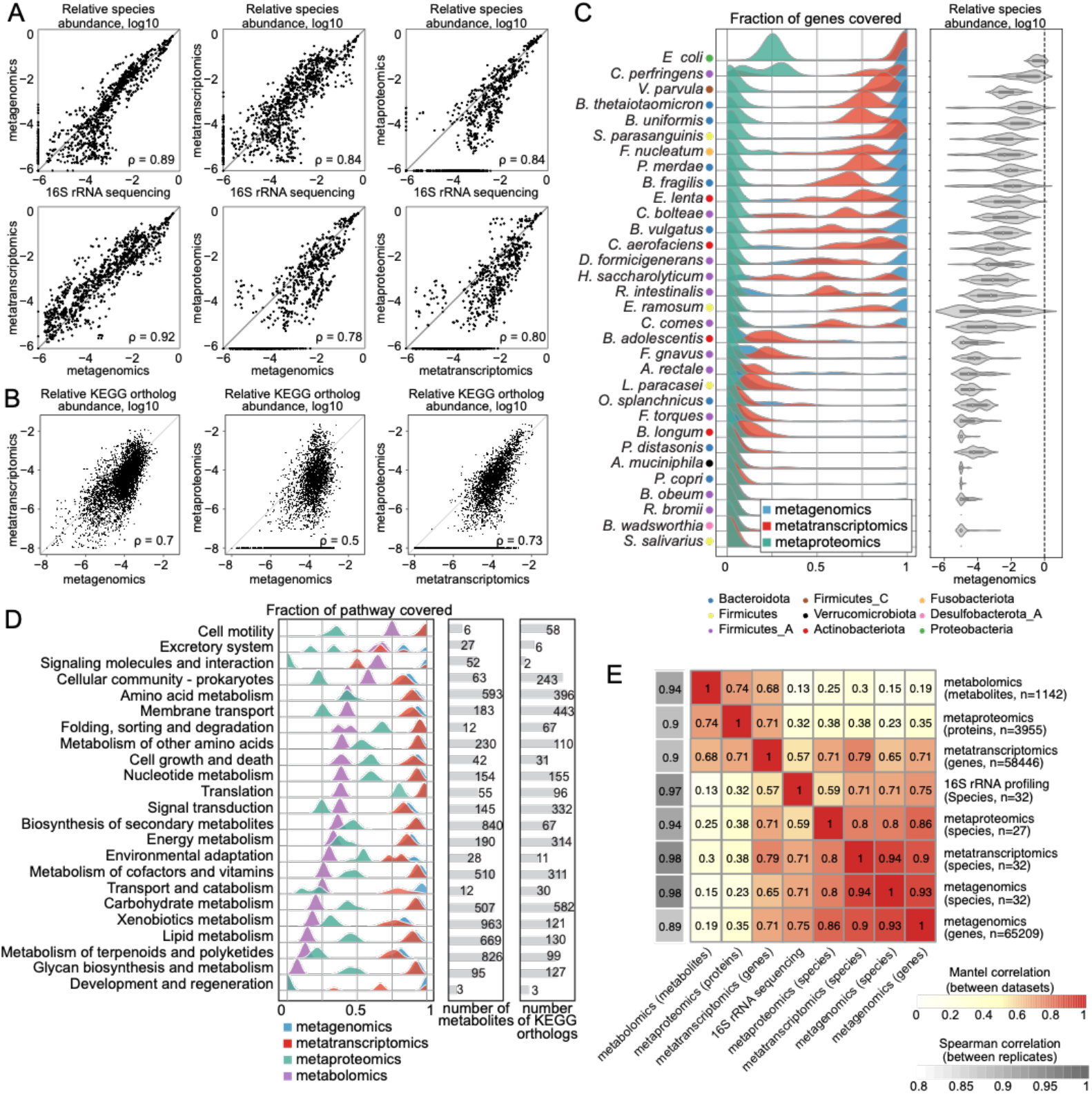
Comparison of species and feature abundances and functional coverage across omics methods. (A) Scatter plots representing species abundance defined as relative abundance of corresponding omics measurement in each sample. Each dot represents single species abundance in one sample. ρ - Spearman correlation coefficient. (B) Scatter plots representing gene, transcript or protein abundance linked through KEGG orthology. Each dot represents a single KEGG ortholog in one sample. (C) Left: Genome coverage of each of the omics datasets across samples for each species, right: relative species abundance estimated by metagenomics. The fraction of coverage is defined as the number of genes to which at least one read was mapped (for metagenomics and metatranscriptomics), or the number of detected proteins for metaproteomics divided by the total number of genes in the corresponding genome. (Metagenomics n = 75 samples, metatranscriptomics n = 101 samples and metaproteomics n = 112 samples) (D) KEGG pathway coverage. For the metabolomics dataset, pathway coverage is defined as the number of unique pathway metabolites detected in at least one sample, divided by the total number of metabolites in the pathway. For metagenomics, metatranscriptomics and metaproteomics, KEGG orthologs are used instead of pathway metabolites. (E) Heatmap of Mantel correlations across omics methods and Spearman correlation between replicates within each omics method.

### Consistency of functional profiles across omics measurements

For each protein-coding gene of each species, we can compare relative abundances across the three molecular layers: gene (metagenomics), transcript (metatranscriptomics), and protein (metaproteomics). We performed such pairwise comparisons both for individual genes across all species (Supplementary Figure 5) and for genes grouped based on KEGG orthology (Kanehisa *et al,* 2017) (Figure 2B). The correlation between metagenomic and metaproteomic estimates of gene and protein abundances was moderate (ρ = 0.5 for KEGG grouped features and ρ = 0.48 for all non-zero genes and proteins). Metatranscriptomics and metaproteomics were the most similar (ρ = 0.73 for KEGG orthologs and ρ = 0.60 for transcripts and proteins), followed by metagenomics and metatranscriptomics (ρ = 0.7 for KEGG orthologs and ρ = 0.61 for genes and transcripts).

To systematically assess how much information on the functional level is captured by metagenomics, metatranscriptomics and metaproteomics for different species, we estimated gene and pathway coverage by calculating the proportion of genes or pathways that were detected by each method (Figure 2C and D). We found that 18 out of 32 species had an almost complete coverage (> 90%) in metagenomics, indicating that for these species most of the genes were recovered in all samples measured in this experiment (Figure 2C; in total 101,559 out of 103,921 possible protein-coding genes were detected at least once in the metagenomics dataset). This was not the case for 14 low abundant species, for which the average gene content coverage was < 20%. For metatranscriptomics, the coverage was generally lower than for metagenomics (91,094 out of 103,921 possible transcripts detected at least once). This is however expected as not all genes are expressed in any given condition. Metaproteomics coverage was found to be much lower than metagenomics and metatranscriptomics (9,144 out of 103,921 predicted proteins). This may be due to the limited dynamic range: In contrast to mass-spectrometry-based measurements, sequencing-based methods include an amplification step that increase the amount of material and makes it possible to cover rare transcripts and genes. For *Escherichia coli,* the most abundant species in our synthetic community, the maximum coverage of proteins across all samples did not exceed 30% (1,428 proteins out of 4,978 (29%) predicted proteins compared to 4,978 genes out of 4,978 predicted genes (100%) for metagenomics and 4,962 transcripts out of 4,978 transcripts (99%) for metatranscriptomics). This result is lower than state-of-the-art single species proteomics experiments where around ~62% (2,586 detected proteins out of 4,189 predicted proteins) of bacterial proteins are captured (Mateus *et al,* 2020), likely due to increased sample complexity in the community context, the increased search space of proteins, and the presence of highly similar protein sequences in homologous proteins (where peptides cannot be unambiguously mapped to one protein).

Since metabolomics data reflects the total pools of metabolites in the sample and cannot be analysed at the species level, we assessed the coverage of metabolic pathways defined in the KEGG database and compared it to pathway coverages by other omics methods (Figure 2D). For metabolic pathways annotated in bacterial genomes, we observed an average pathway coverage of 35% for metabolomics, as compared to 44% for metaproteomics and 86% for metatranscriptomics. Even though direct comparison of both methods is challenging, we believe that the lower coverage for metabolomics has several explanations. First, we measured metabolites in supernatant samples, where rich medium components mask the signal (e.g. amino acids, peptides and polysaccharides), and extracellular products of bacterial metabolism, especially produced by only one or few species, may therefore not be detected. Second, only a subset of all metabolites present in the bacterial cell will be secreted outside of the cell. Third, to calculate metabolic pathway coverage, we assumed that each pathway consists of metabolites that are produced or consumed by metabolic enzymes annotated in bacterial genomes, which is likely an overestimation of pathway sizes, since presence of an enzyme in the genome does not necessarily imply that this enzyme is expressed, and that the corresponding metabolite will be produced and measured extracellularly.

To further compare the samples measured with different omics methods, we performed a Mantel test, which measures a correlation coefficient between sample similarity matrices calculated based on each omics data type individually (Figure 2E). For example, while it is not possible to directly compare matrices of species and protein abundances, it is possible to calculate sample similarity matrices for these two methods that can then be compared with each other. Hierarchical clustering of Mantel correlation coefficients revealed two clusters: one containing species abundance data (from metagenomics, metatranscriptomics, and metaproteomics) and gene abundance (metagenomics); and a second cluster containing transcript abundance data (metatranscriptomics), protein abundance data (metaproteomics), and metabolite abundances. The emergence of these clusters can be explained by the nature of data used to calculate sample distance matrices: species and gene abundances in one cluster, and functional feature abundances in the other cluster. Notably, transcript abundance as measured by metatranscriptomics showed a high correlation (≥0.57) with sample distance matrices of all other omics measurements, underlining that this method captures both species abundance and functional information in our experiment. Altogether, metatranscriptomics was found to be the most universal and versatile readout, as it can both provide robust and sensitive estimates of species abundance, and at the same time reflects functional changes, which are in concordance with protein changes detected by metaproteomics.

### Chlorpromazine treatment strongly affects community composition

After testing the technical consistency between omics measurements in a synthetic microbial community, we explored the impact of drug perturbations on the community composition and the respective responses at species, gene, transcript, protein and metabolite levels. For the control condition and all perturbations (chlorpromazine, metformin and niclosamide), similar dynamic changes in alpha diversity were observed over time. In general, the alpha diversity (inverse Simpson index) increased as the community grew over time after inoculation, however, this increase was lower for chlorpromazine compared to the other drugs and the control condition (Supplementary Figure 6A, B). We observed different community dynamics between runs A and B during the exponential phase: *E. coli* and *C. perfringens* were the most abundant species in all conditions in run A (Fig. 3A, Supplementary Figure 6C), while *E. coli* dominated community composition during exponential phase in run B. However, community compositions became more similar between the runs at 43 h after drug treatment (Supplementary Figure 7). These analyses revealed that the addition of metformin and niclosamide had negligible effects on the community composition, while chlorpromazine treatment shifted the community composition in both runs.

**Figure 3.**
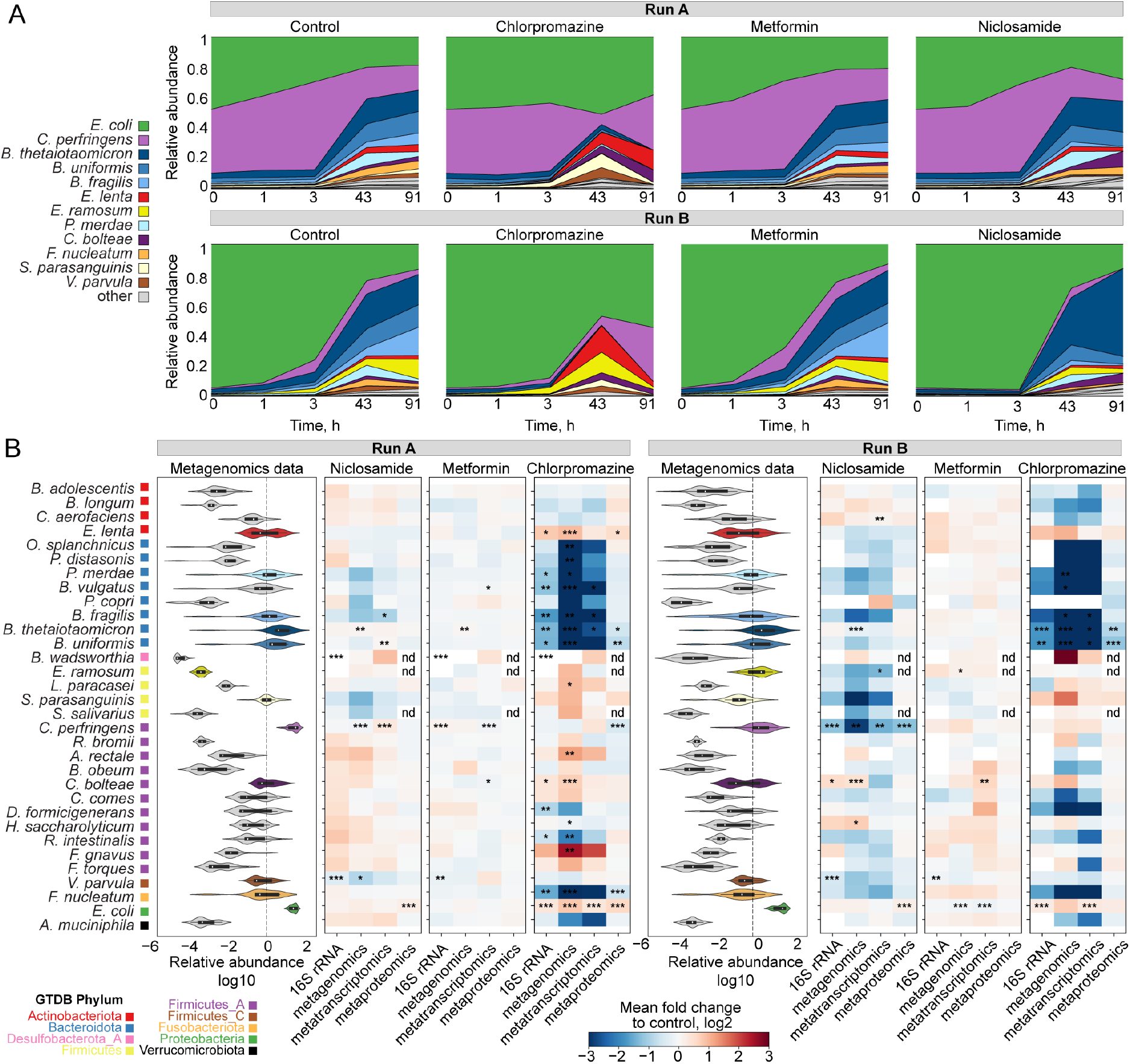
Changes in community composition upon drug perturbation. (A). Relative species abundance changes over time in the three drug conditions and control. Time 0 indicates timepoint of the drug addition 5 h after the passage in the fresh medium. Relative abundance measured from metagenomics data. (B). Left, distribution of relative species abundance for each species across all samples (all conditions and timepoints). Right, heatmap of species abundance fold changes measured by different omic methods for each drug condition versus control. Significance of changes estimated by the ANCOM test is indicated by asterisks: * changes detected at 0.7 threshold of W statistic; ** changes detected at 0.8 threshold; *** changes detected at 0.9 threshold; nd – not detected.

To identify differentially abundant species after drug perturbation, we analysed the composition of microbiomes by comparing species abundances in drug-treated samples against control samples estimated by each omics type (Figure 3B) (ANCOM (Mandal *et al*, 2015)). This analysis revealed that most members of the Bacteroidota phylum *(Odoribacter splanchnicus, Parabacteroides distasonis, Phocaeicola vulgatus, Bacteroides fragilis, Bacteroides thetaiotaomicron* and *Bacteroides uniformis*) were less abundant in chlorpromazine-treated samples. This reduction in Bacteroidota abundance was detected across all four omics methods capturing community composition, indicating that each of these methods is capable of detecting strong signals of species abundance change. In addition to Bacteroidota, *Fusobacterium nucleatum* was found to be less abundant in chlorpromazine-treated samples. In contrast, the other two drugs did not cause major shifts in relative abundances: although ANCOM test identified significant changes of abundance of several species, their relative abundance was not changing more than two-fold (Figure 3B). In summary, we found a consistent and substantial depletion of species belonging to the phylum Bacteroidota upon chlorpromazine treatment.

### Multi-omics measurements capture functional response of the community to all three drugs

As compositional shifts do not provide information on the mechanisms of response of each community member, we investigated these functional responses in more detail by performing differential analysis of metatranscriptomic, metaproteomic and metabolomic datasets after a normalization step wherein taxonomic abundance effects were reduced (see “Gene, transcript and protein counting” in the Methods section). The highest number of differentially abundant transcripts, proteins and metabolites were found in samples treated with chlorpromazine (adjusted p-value < 0.001 and absolute fold change > 4 compared to control for metatranscriptomics, adjusted p-value < 0.05 and absolute fold change > 1.5 for metaproteomics and metabolomics; Figure 4A), which is in line with our findings that chlorpromazine caused the largest disruption to bacterial community (Figure 3B). Transcriptional response to chlorpromazine is detected already after 15 minutes of treatment across species belonging to different phyla, suggesting that, although Bacteroidota show the strongest response, other species also adapt their gene expression.

**Figure 4.**
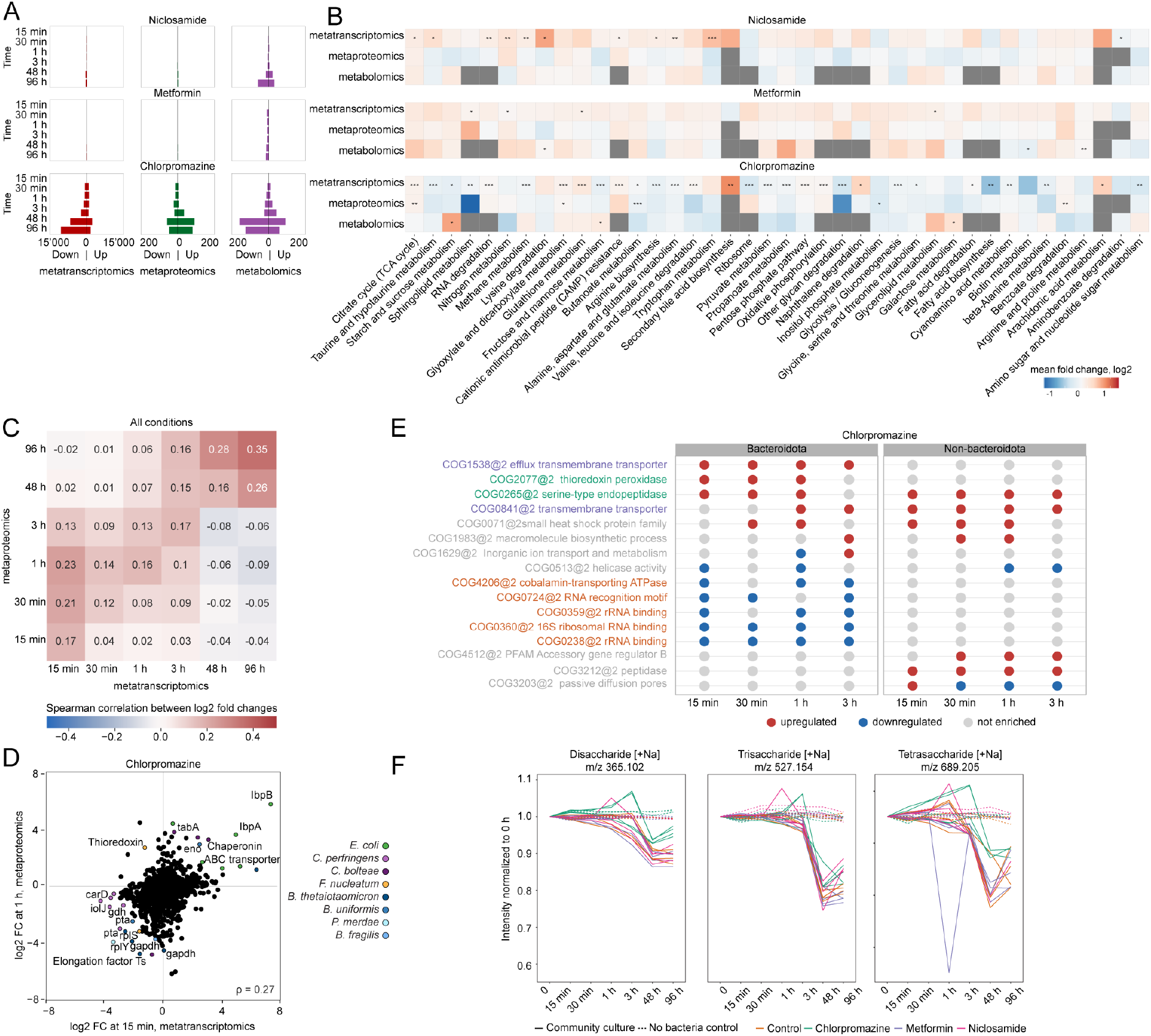
Functional analysis of transcript, protein and metabolite response after niclosamide, metformin or chlorpromazine treatment. A) Number of differentially abundant transcripts, proteins and metabolites. B) Pathway enrichment analysis across all conditions and time points. C) Heatmap representing Spearman correlation between fold changes (relative to control) detected by metatranscriptomics and metaproteomics across all drug perturbations. D) Scatterplot depicting protein fold changes (relative to control) detected after 15 min of chlorpromazine exposure by metatranscriptomics versus after 1 h of exposure by metaproteomics. E) COG enrichment analysis differentiating between species susceptible to chlorpromazine treatment (Bacteroidota) and non-susceptible species (non-Bacteroidota). COGs that are enriched in upregulated genes are coloured in red, while COGs that are enriched in downregulated genes are coloured in blue. Only COGs that were found to be significantly enriched in at least three out of four early time points are shown. COG names that are coloured are discussed in more detail in the main text. F) Di-, tri- and tetra-saccharide abundances as measured by untargeted metabolomics (metabolite annotation is based on m/z values indicated in the panel titles). The lines are coloured according to the experimental conditions (chlorpromazine, metformin, niclosamide and control), and the line type represents whether these are community culture or non-bacterial controls.

In order to evaluate similarities between functional responses across omics data types, we performed pathway enrichment analysis of differentially abundant features between drug treatment and controls across all time points using KEGG pathway annotations (Figure 4B). In general, we detected less overlap between omics layers on the functional level compared to species abundance analysis, as no single pathway was statistically significant in the enrichment analysis of all three functional omics datasets. Across all conditions, five pathways were found to be significantly enriched upon drug treatment compared to the control condition in two omics data types, while 35 pathways were statistically significantly enriched in only one omics dataset. The largest number of significantly enriched pathways was found in chlorpromazine-treated samples for metatranscriptomics data.

Several pathways were significantly overrepresented (pFDR<0.001 for metatranscriptomics and pFDR<0.05 for metaproteomics and metabolomics) within the set of up- and downregulated features (transcripts/proteins/metabolites) in metformin-treated samples. For example, three pathways, i) lysine degradation, ii) biotin metabolism and iii) arginine and proline metabolism, were enriched in differentially abundant metabolites. Further inspection of metabolites involved in these pathways showed that their abundance also decreased upon addition of metformin in the non-bacterial control samples (Supplementary Figure 8). This indicates that metformin primarily interferes with the measurement of these metabolites, probably due to their chemical similarity, underlining the importance of including non-bacterial control samples to study drug response. However, we cannot exclude that metformin also interacts with lysine and arginine metabolism pathways in bacteria, as reported before (Pryor *et al,* 2019; Forslund *et al,* 2015). In general, we did not observe substantial effects of metformin neither on community composition nor on transcript or protein abundance in our study, at least at the concentrations used. In previous experiments, metformin at the same concentration had an effect on several of the tested species grown in single culture (Maier *et al,* 2018), however it is possible that these species show a different behaviour in a community setting (D’hoe *et al,* 2018).

For niclosamide-treated samples, ten pathways were significantly enriched (pFDR<0.001) among regulated transcripts, including amino acid and nitrogen metabolism. Transcripts of nitrogen metabolism pathway upregulated in the early time points (15 min, 30 min, 1 h, 3 h) were annotated as NAD-specific glutamate dehydrogenase (belonging to the Cluster of Orthologous Groups COG0334 from the EggNOG database present in *B. thetaiotaomicron, P. vulgatus, B. fragilis*), hydroxylamine reductase (COG1151 in *C. perfringens, B. uniformis*) and carbamate kinase (COG0549 in *Eggerthella lenta*) (Supplementary Figure 9). Previously, NAD-specific glutamate dehydrogenase was found to be upregulated in response to nitrogen availability in *Mycobacterium smegmatis,* where it is assumed to have a de-aminating activity (Harper *et al,* 2010). Furthermore, hydroxylamine reductase and carbamate kinase are enzymes belonging to the family of oxidoreductases which both act on nitrogenous compounds. Therefore, the upregulated pathway and its transcripts suggest an increased metabolization of nitrogen in niclosamide-treated samples. Further examination of our metabolomic dataset revealed that niclosamide gets degraded in both runs of the experiment (Supplementary Figure 10). Additional follow-up experiments are needed to elucidate the mechanisms underlying the microbial degradation of niclosamide and the roles of individual community members.

### Chlorpromazine induces stress response and metabolic changes in the community

Since the number of differentially abundant features and pathways was high in chlorpromazine-treated samples (Figure 4A and 4B), we tested whether there are features that change concordantly across omics layers. We first compared transcript and protein fold changes upon perturbation, which revealed general agreement between relative changes in gene expression and protein abundance, with transcript fold changes at each time point correlating more strongly with protein changes at later time points (Figure 4C, Supplementary Figure 11), likely reflecting the delay between transcription and translation processes. Based on this analysis, we assessed the most prominent and concordant changes between metatranscriptomics and metaproteomics 15 min and 1 h after chlorpromazine addition, respectively (Figure 4D). Consistent with the observed relative species abundance changes, the most concordantly downregulated features were proteins and genes of Bacteroidota species and *F. nucleatum,* including ribosomal proteins, elongation factors, and central carbon metabolism enzymes *gldA* (glycerol dehydrogenase), *gapdh* (glyceraldehyde 3-phosphate dehydrogenase), and *pta* (phosphate acetyltransferase), the latter two being downregulated in several species (Figure 4D). Furthermore, the most upregulated features found both in metatranscriptomics and metaproteomics were stress response genes in *E. coli,* such as the small heat shock proteins IbpA and IbpB (Inclusion body-associated protein A and B), other chaperones, and ABC transporters. IbpA and IbpB serve as a first line of defence against protein aggregation (Miwa *et al,* 2021). In addition to *ibpA* and *ibpB,* we found upregulation of the transcriptional regulator *rpoH* and the chaperones *dnaK* and *groEL,* which are also involved in heat shock response (Yura, 2019) (Supplementary Figure 12). Together, these results show that chlorpromazine causes the activation of a stress response in *E. coli,* probably due to induction of protein aggregation either directly or indirectly.

We then tested whether genes associated with stress response were differently regulated between chlorpromazine-susceptible and non-susceptible species. Two COGs related to the stress response were enriched in upregulated genes in at least two of the four early time points in the depleted (susceptible) species (Figure 4E, annotated in green, Supplementary Figure 13). One of them, COG0265, is upregulated by both susceptible and non-susceptible species and encompasses serine proteases (e.g., HtrA proteins such as DegP and DegQ), which represent an important class of chaperones and heat-shock-induced serine proteases, protecting periplasmic proteins. Furthermore, two COGs enriched in upregulated genes were related to (multidrug) transporter activity. COG1538, which contains genes annotated as membrane protein OprM, was the only COG enriched in upregulated genes by Bacteroidota in all four early time points (Fig 4F, annotated in purple). In *Pseudomonas aeruginosa,* OprM is part of MexAB-OprM, a multidrug efflux pump of the resistance-nodulation-cell division (RND) superfamily, where it plays a central role in multidrug resistance by transporting drugs from the cytoplasm across the inner and outer membranes outside the cell envelope (Tsutsumi *et al,* 2019; Alekshun & Levy, 2007). RND-efflux pumps are found in a number of Gram-negative bacteria, for example, AcrAB-TolC is found in *E. coli* (Du *et al,* 2018) while *Bacteroides fragilis* harbours multiple copies of RND pumps BmeABC (Ghotaslou *et al,* 2018). Further, in addition to COG1538 (OprM homologues), also COG0841 containing homologues of the MexB/AcrB/bmeB protein (Figure 4E, also annotated in purple) was found to be enriched in upregulated genes, both in Bacteroidota and non-Bacteroidota species. These observations suggest an important role of the AcrAB-TolC/MexAB-OprM/bmeABC efflux pumps in determining chlorpromazine susceptibility. Indeed, a recent study showed that chlorpromazine is both a substrate and an inhibitor of the AcrB multidrug efflux pump in *Salmonella enterica* and *E. coli* (Grimsey *et al,* 2020). Together, our results suggest that chlorpromazine could also be an inhibitor of BmeB, the AcrB/MexB homologue in Bacteroidota species and that this, potentially in combination with protein aggregation, could be one of the reasons explaining why Bacteroidota are affected by chlorpromazine treatment.

Finally, depletion of Bacteroidota and downregulation of their genes involved in polysaccharide uptake might explain the enrichment of “Starch and sucrose” and “Fructose and mannose metabolism” pathways among metabolites increased upon chlorpromazine treatment (Figure 4B). As *Bacteroides* species are known to be capable of metabolizing a wide variety of polysaccharides (Schwalm & Groisman, 2017), we believe that the higher abundance of polysaccharides after chlorpromazine treatment (Figure 4F) measured by metabolomics is a result of their reduced consumption by these species.

Taken together, by integrating multi-omics measurements, we propose that a series of events happens upon treatment by chlorpromazine i) a stress response is induced across several bacterial species with overexpression of *ibpA* and *ipbB* chaperones being the most pronounced response in *E. coli*; ii) this stress response involves upregulation of AcrB/BmeB type of RND pumps, which may be bound and blocked by chlorpromazine in a species-specific manner; iii) Bacteroidota species are more susceptible to chlorpromazine and are quickly depleted from the community, which results in iv) increase in polysaccharide levels in the culture medium due to the inability of the remaining community members to utilize them.

## Discussion

In this study we evaluated the impact of drug perturbations on a synthetic gut microbial community by analysing five different omics data types in a highly controlled *in vitro* experiment. In general, we found concordance between all omics data types regarding the estimation of community composition (taxonomic profiling). To our knowledge, this is the first study to systematically compare taxonomic profiles obtained by four omics data types and can thus serve as a baseline for integrating different data types in “in natura” settings. Using the synthetic community, we could show a high correlation between metagenomics and metatranscriptomics (ρ = 0.92), similar to a previous study that used only these two omics methods (ρ = 0.81, (Heintz-Buschart & Wilmes, 2018)). The taxonomic profiles obtained from our metaproteomics dataset, which is increasingly used in microbiome studies (e.g. Kleiner *et al,* 2017; Kleikamp *et al,* 2021), showed correlations between ρ = 0.78 and ρ = 0.84 with all other omics for species with a relative abundance higher than 1%. Although the number of detected proteins and the detection limits remain to be improved, we showed that species abundance estimates can be derived from metaproteomics in a relatively simple, defined microbial community.

Of the three drugs used for perturbation, only chlorpromazine caused a large disturbance in the community composition. Surprisingly, metformin, which has been shown to alter the gut microbiome in patients (Forslund *et al*, 2015; Wu *et al*, 2017), did not perturb the community in our study, even though our earlier study suggested that the growth of at least four different species is inhibited by metformin at the concentration used in monocultures (*F. nucleatum, B. longum, P. copri* and *P. merdae,* (Maier *et al,* 2018)). This observation hints at a protective effect from the community, although this protective effect is not caused by drug degradation, as metformin concentrations remained high during the course of experiment (Supplementary Figure 10). Similarly, niclosamide was expected to cause a depletion of most of the members of the synthetic community, except for *E. coli* and *B. wadsworthia* (Maier et al. 2018), which was not observed in this study, also pointing to community-related protection effects. Our metatranscriptomic data revealed an upregulation of genes related to nitrogen metabolism, while niclosamide concentration decreased during incubation, which was not observed in the non-bacterial controls. Therefore, we believe that certain species are capable of degrading niclosamide, which ultimately protected the whole community against possible inhibitory effects of niclosamide treatment.

For chlorpromazine, the observed depletion of Bacteroidota species was in concordance with single species experiments (Maier *et al,* 2018). The antibiotic activity of chlorpromazine was reported relatively soon after its first usage in the nineteen-fifties (Dinan & Cryan, 2018; Kristiansen & Vergmann, 1986). Its antibiotic mechanism of action is described to be multifold and includes effects on the cell membrane, energy generation and interference with cell replication due to DNA intercalation in *E. coli* (Grimsey *et al,* 2020). In our study, several genes and proteins related to protein aggregation were upregulated in metatranscriptomic and metaproteomic datasets in *E. coli* and other community members. One study already reported protein aggregation of bovine insulin after chlorpromazine treatment (Bhattacharyya & Das, 2001). However, it remains unclear whether chlorpromazine can cause protein aggregation in microbes either directly or indirectly, a hypothesis that should be followed-up in future experiments.

Finally, we identified upregulation of RND-type efflux pumps in the Gram-negative bacteria, even in the Bacteroidota species that were severely depleted. It was recently shown that in *S. enterica* and *E. coli,* chlorpromazine is both a substrate and an inhibitor of AcrB, the inner membrane transporter of the tripartite system AcrAB-TolC, which is an RND-type efflux pump (Grimsey *et al,* 2020; Bailey *et al,* 2008a). Based on our data, we hypothesise that BmeB, the AcrB homologue in Bacteroidota, is also susceptible to chlorpromazine inhibition as we found upregulation of this and related genes, similar to what has been described by others in single species experiments (Grimsey *et al*, 2020). The suggested mechanism could be of significance in the battle against the rising multidrug resistance of *Bacteroides fragilis,* a commensal bacterium that can act as a virulent pathogen when it escapes its normal niche (Wexler, 2007, 2012; Niestępski *et al,* 2019). However, chlorpromazine’s antimicrobial activity generally occurs at concentrations higher than those clinically achievable (Grimsey & Piddock, 2019). Therefore it is possible that, similarly as suggested for *S. enterica,* chlorpromazine could act as an antimicrobial adjuvant for Bacteroidota where its inhibition of RND-type efflux pumps prevents the extrusion of administered antibiotics (Grimsey *et al,* 2020). From the perspective of human health, these results underline the detrimental effect of antipsychotics on the gut microbiome reported before (Dinan & Cryan, 2018). However, the revealed phylum-specific differences provide an opportunity to explore whether complementation of antipsychotic therapy with Bacteroidota-promoting dietary interventions could improve mental health and increase patients’ quality of life by restoring a healthy microbiota (Patnode *et al*, 2019).

In conclusion, we directly compared data from multiple omics methods and showed that they agree on species abundance estimation of a defined and drug-perturbed microbial community *in vitro*. Those methods that are able to detect functional information also correlate with each other, albeit to a lower degree. We could also confirm expected time delays between transcriptional and translational responses to perturbations, underlining that these methods reveal biological insights that happen at different time scales. Although multi-omics analysis of natural communities is hampered by their increasing complexity, combining multiple omics measurements allows to measure the response of the community to perturbations across molecular layers and provides information that is not achievable by any method alone.

## Methods

### Species and drug selection

The species used in this study represent a subset of abundant and prevalent species from the human gut. In total, 32 species were selected based on our previous work (Tramontano *et al,* 2018; Maier *et al,* 2018). The bacterial isolates were received from DSMZ, BEI Resources or ATCC and Dupont Health & Nutrition. The drugs were chosen because of their antimicrobial activity (Maier *et al*, 2018) and diversity in therapeutic usage.

### Reference genomes

Reference genomes were downloaded from RefSeq on March 2019 (release 92) and reannotated using Prokka v1.14.0 (Seemann, 2014). Taxonomic classification was based on GTDB taxonomy release 95 (Parks *et al*, 2018) and inferred using GTDB-Tk v1.3.0 (Chaumeil *et al*, 2020; Matsen *et al*, 2010; Jain *et al*, 2018; Hyatt *et al*, 2010; Price *et al*, 2010; Eddy, 2011; Ondov *et al*, 2016). Further functional annotations (e.g. KEGG orthology and eggNOG orthologous group) were retrieved using eggNOG-mapper v2.0.1 which is based on eggNOG v5.0 (Huerta-Cepas *et al*, 2017). A cladogram was built by pruning the species cladogram from GTDB (bac120.tree, release 95) using the ETE toolkit (Huerta-Cepas *et al,* 2016).

### Medium and drug preparation

mGAM medium was prepared according to manufacturer’s instructions (HyServe GmbH & Co.KG, Germany, produced by Nissui Pharmaceuticals) and all the single species were grown in this medium except *V. parvula* (Todd-Hewitt Broth (Sigma-Aldrich) + 0.6% sodium lactate) and *B. wadsworthia* (mGAM + 60 mM sodium formate + 10 mM taurine). All media were placed in anaerobic chamber 1 day before use under anoxic conditions (Coy Laboratory Products Inc.) (2% H2, 12% CO2, rest N2). Chlorpromazine (TCI Chemicals) and niclosamide (Santa Cruz Biotechnology) were added from DMSO stock solution. Metformin (Sigma) was added as powder directly into the medium after which the medium was filter-sterilized. Final concentrations of each drug were chosen based on previous work (Maier et al. 2018) with concentrations of 5 mM for metformin and 20 μM for chlorpromazine and niclosamide. The higher concentration for metformin is motivated by previously published data, which showed that a concentration of 20 μM was not sufficient to impair growth of gut microbiome members *in vitro* ((Maier *et al,* 2018); Extended Data Figure 4).

### Experimental set-up and sample collection

Species were pre-inoculated in isolation on liquid mGAM medium from pure stocks and incubated at 37 °C under anaerobic conditions for a period of 3 or 5 days, depending on the growth rate of each species (see Figure 1). The monocultures were subsequently mixed in equal proportions based on their optical density (OD) and then inoculated in 100 mL of mGAM liquid medium. To allow species to reach a stable state (stabilization phase), the mixed culture was grown for 48 hours after which 1 mL was transferred to fresh medium. In total, 3 passages were performed and after the second transfer OD measurements were taken to determine the start of the exponential phase.

Following the stabilization phase, the mixed community was inoculated in medium prepared with one single drug or DMSO (control) as soon as the community reached the exponential phase (OD roughly equal to 2-3). The cultures were subsequently sampled (3 mL) at fixed time intervals (0 minutes, 15 minutes, 30 minutes, 1 hour, 3 hours, 48 hours), transferred to fresh medium (with drugs or DMSO) after 48 hours and then sampled again 48 hours later (or 96 hours after the start of the experiment). The whole experiment was performed twice (labelled as run A and run B).

1.5 mL of each collected sample was centrifuged (30 seconds at max speed) after which the supernatant was removed and the cell pellet was stored at −80 °C until further processing for DNA and RNA extraction. For protein and metabolite extraction, again 1 mL of each collected sample was centrifuged (30 seconds at max speed) and 450 μL of supernatant was used for metabolite extraction or protein extraction (secreted proteins) while the cell pellet was used for protein extraction (proteins in the cells). The remainder of the samples was frozen at −80 °C as backup.

### DNA and RNA extraction

Genomic DNA and total RNA were extracted from the same flash-frozen samples using Allprep Powerfecal DNA/RNA kit (Qiagen, Hilden Germany) following the manufacturer’s protocol but an additional phenol-chloroform extraction step of 700 μL was performed after lysis. DNA yield was measured by using Qubit™ dsDNA HS Assay Kit (Qubit, Waltham, Massachusetts, USA), split into two aliquots for ribosomal 16S rRNA amplicon sequencing and metagenomic shotgun sequencing and was stored at −20°C. RNA yield was measured via Bioanalyzer (Agilent, Santa Clara, California, USA) with Pico and Nano chips depending on the sample concentration and stored at −80°C for further analysis.

### 16S rRNA amplicon, metagenomic and metatranscriptomic sequencing

For 16S rRNA amplicon sequencing, extracted DNA was amplified using primers targeting the V4 region of the 16S rRNA gene on the F515 and R806 primer pair (Caporaso *et al*, 2011). PCR was performed according to the manufacturer’s instructions of the KAPA HiFi HotStart PCR Kits (Roche, Basel Switzerland) using barcoded primers and a two-step PCR protocol (NEXTflex™ 16S V4 Amplicon-Seq Kit, Bioo Scientific, Austin, Texas, USA). PCR products were pooled and purified using size-selective SPRIselect magnetic beads (0.8 left-sized, Beckman Coulter, Brea, CA, USA). The library was then diluted to 6pM for sequencing. The library was sequenced on an Illumina (San Diego, USA) MiSeq platform using 2 x 250 bp paired-end reads at Genomics Core Facility (European Molecular Biology Laboratory (EMBL), Heidelberg, Germany).

Metagenomic libraries for all samples were prepared using the NEB Ultra II and SPRI HD kits with a targeted insert size of 350, and sequenced on an Illumina HiSeq 4000 platform (Illumina, San Diego, CA, USA) in 2×150bp paired-end with the aim of 1.5 Gbp average setup at the Genomics Core Facility (EMBL, Heidelberg, Germany).

RNA samples were depleted for ribosomal RNA using the NEBNext Bacteria rRNA Depletion Kit (New England Biolabs, Ipswich, Massachusetts, USA). Samples were pooled into a library using the NEBNext Ultra II Directional RNA Library Prep Kit (New England Biolabs) and subsequently sequenced on Illumina NextSeq500 platform (75 bp; single end) at Genomics Core Facility (EMBL, Heidelberg, Germany).

Quality control of raw reads was performed using NGLess (Coelho *et al,* 2019). For metagenomics, reads were trimmed to the longest subread where each base had a Phred score of at least 25. For metatranscriptomics, a sliding window approach was used and reads were trimmed to the longest subread with an average Phred score of 20 (window size: 4 bp). Resulting reads shorter than 45 bp were discarded. To remove possible human contamination, all reads were mapped against a human reference database (release GRCh38.p10, Ensembl (Zerbino *et al,* 2018)) using NGLess and samtools (Li *et al,* 2009). Reads with an identity threshold >= 90% were discarded. For metatranscriptomics specifically, rRNA reads were also removed from the dataset using SortMeRNA (Kopylova *et al,* 2012) with default parameters.

### Protein extraction

Sample preparation, including protein extraction, digestion and peptide purification was performed according to the in-StageTip protocol (Kulak *et al,* 2014, 20) with automation on an Agilent Bravo liquid handling platform according to (Geyer *et al,* 2016). In brief, samples were incubated in the PreOmics lysis buffer (P.O. 00001, PreOmics GmbH) for reduction of disulfide bridges, cysteine alkylation and protein denaturation at 95°C for 10 min. Samples were sonicated using a Bioruptor Plus from Diagenode (15 cycles of 30 s). The protein concentration was measured using a tryptophan assay. In total, 200 μg protein of each organism were further processed on the Agilent Bravo liquid handling platform by adding trypsin and LysC (1:100 ratio - μg of enzyme to μg of sample protein). Digestion was performed at 37 °C for 4 h.

The peptides were purified in consecutive steps according to the PreOmics iST protocol (www.preomics.com). After elution from the solid phase extraction material, the peptides were completely dried using a SpeedVac centrifuge at 60°C (Eppendorf, Concentrator plus). Peptides were suspended in buffer A* (2% acetonitrile (v/v), 0.1% trifluoroacetic acid (v/v)) and sonicated for 30 min (Branson Ultrasonics, Ultrasonic Cleaner Model 2510).

### Metaproteomics

Samples were analyzed using a liquid chromatography (LC) system coupled to a mass spectrometer (MS). The LC was an EASY-nLC 1200 ultra-high pressure system (Thermo Fisher Scientific) and was coupled to a Q Exactive HFX Orbitrap mass spectrometer (Thermo Fisher Scientific) using a nano-electrospray ion source (Thermo Fisher Scientific). Purified peptides were separated on 50 cm HPLC-columns (ID: 75 μm; in-house packed into the tip with ReproSil-Pur C18-AQ 1.9 μm resin (Dr. Maisch GmbH)). For each LC-MS/MS analysis about 500 ng peptides were separated on 100 min gradients.

Peptides were separated with a two-buffer-system consisting of buffer A (0.1% (v/v) formic acid) and buffer B (0.1% (v/v) formic acid, 80% (v/v) acetonitrile). Peptides were eluted with a linear 70 min gradient of 2-24% buffer B, followed stepwise by a 21 min increase to 40% buffer B, a 4 min increase to 98% buffer B and a 5 min wash of 98% buffer B. The flow rate was constant at 350 nl/min. The temperature of the column was kept at 60°C by an in-house-developed oven containing an Peltier element, and parameters were monitored in real time by the SprayQC software (Scheltema & Mann, 2012).

First, data dependent acquisition (DDA) was performed of each single organism to establish a library for the data independent acquisition (DIA) of the community culture samples. The DDA scans consisted of a Top15 MS/MS scan method. Target values for the full scan MS spectra were 3e6 charges in the 300–1650 m/z range with a maximum injection time of 25 ms and a resolution of 60,000 at m/z 200. Fragmentation of precursor ions was performed by higher-energy C-trap dissociation (HCD) with a normalized collision energy of 27 eV. MS/MS scans were performed at a resolution of 15,000 at m/z 200 with an ion target value of 5e4 and a maximum injection time of 120 ms. Dynamic exclusion was set to 30 s to avoid repeated sequencing of identical peptides.

MS data for the community culture samples were acquired with the DIA scan mode. Full MS scans were acquired in the range of m/z 300–1650 at a resolution of 60,000 at m/z 200 and the automatic gain control (AGC) set to 3e6. The full MS scan was followed by 32 MS/MS windows per cycle in the range of m/z 300-1650 at a resolution of 30,000 at m/z 200. A higher-energy collisional dissociation MS/MS scans was acquired with a stepped normalized collision energy of 25/27.5/30 eV and ions were accumulated to reach an AGC target value of 3e6 or for a maximum of 54 ms.

The MS data of the single organisms and of the community cultures were used to generate a DDA-library and the direct-DIA-library, respectively, which were computationally merged into a hybrid library using the Spectronaut software (Biognosys AG). All searches were performed against a merged protein FASTA file of our reference genomes annotated using Prokka (see above). Searches used carbamidomethylation as fixed modification and acetylation of the protein N-terminus and oxidation of methionines as variable modifications. Trypsin/P proteolytic cleavage rule was used, permitting a maximum of 2 missed cleavages and a minimum peptide length of 7 amino acids. The Q-value cutoffs for both library generation and DIA analyses were set to 0.01.

### Metabolomics measurements

Untargeted metabolomics analysis was performed as described previously (Fuhrer *et al*, 2011). Briefly, samples were analyzed on a LC/MS platform consisting of a Thermo Scientific Ultimate 3000 liquid chromatography system with autosampler temperature set to 10° C coupled to a Thermo Scientific Q-Exactive Plus Fourier transform mass spectrometer equipped with a heated electrospray ion source and operated in negative or positive ionization mode. The isocratic flow rate was 150 μL/min of mobile phase consisting of 60:40% (v/v) isopropanol:water buffered with 1 mM ammonium fluoride at pH 9 for negative ionization mode or 60:40% (v/v) methanol:water buffered with 0.1% formic acid at pH 2 for positive ionization mode, in both cases containing 10 nM taurocholic acid and 20 nM homotaurine as lock masses. Mass spectra were recorded in profile mode from 50 to 1,000 m/z with the following instrument settings: sheath gas, 35 a.u.; aux gas, 10 a.u.; aux gas heater, 200° C; sweep gas, 1 a.u.; spray voltage, −3 kV (negative mode) or 4 kV (positive mode); capillary temperature, 250° C; S-lens RF level, 50 a.u; resolution, 70k @ 200 m/z; AGC target, 3×10^6^ ions, max. inject time, 120 ms; acquisition duration, 60 s. Spectral data processing including peak detection and alignment was performed using an automated pipeline in R analogous to previously published pipelines (Fuhrer *et al,* 2011). Detected ions were tentatively annotated as metabolites based on accurate mass within a dynamic tolerance depending on local instrument resolving power ranging from 1 mDa at m/z = 50 to 5 mDa at m/z = 1,000 using the Human Metabolome database (Wishart *et al,* 2018) as reference considering [M-H] and [M-2H] ions in negative mode or [M+], [M+H], [M+Na] and [M+K] ions in positive mode and up to two ^12^C to ^13^C substitutions. Of note, this approach precludes the resolution of isomers, of metabolites mapping to the same ion using different adduct assumptions, of unaccounted neutral gains or losses, or of metabolites with slightly distinct masses that nevertheless map to the same ion within the respective local matching tolerance.

### Metabolomics data analysis

Raw intensity values were quantile-normalized separately for ions acquired in positive and negative modes. For further analysis, the data from the two acquisition polarity modes were combined in one table and filtered as follows: only annotated ions were retained; ions annotated to ^13^C-compounds only were removed; for each metabolite, only the ion with the annotation considered most likely was retained (either the ion with highest correlation with the total ion current, or the ion with the largest mean intensity across samples).

### Gene, transcript and protein counting

Metagenomic and metatranscriptomic reads were mapped against a database of reference genomes containing only the species used in this study, using NGLess and samtools, with a minimum match size of 45 and minimum identity of 97. Abundance estimates were produced by counting the number of reads mapping to each genome included in the study. If a read mapped to multiple genes, the count was distributed to each of the genes (e.g. if a read maps to gene X and gene Y, gene X and gene Y each get a count of 0.5).

Proteins quantification and filtering. Proteins were filtered based on the information from the DDA experiment on which peptides are detected in which single species. Metaproteomics report with protein and peptide quantification obtained from Spectronaut software applied to DIA samples was used as input. For each peptide in the community peptide report file, number of exact protein and species matches was calculated. For each protein, only unique peptides that match to one species were left for quantification. For each protein, the peptides were sorted according to the number of samples in which they were detected. Protein abundance was calculated as the mean of three most commonly measured peptides as suggested before. If the number of peptides was less than three, the protein was discarded.

To reduce taxonomic abundance effects in downstream analyses, taxon-specific scaling was performed on metagenomics, metatranscriptomics and metaproteomics as described by (Klingenberg & Meinicke, 2017).

### Species abundance estimation

Multiple computational strategies were used to estimate species abundance. Unless stated otherwise, for all analyses the species abundances resulting from read mapping were used. For this approach, first a database of 16S rRNA regions was constructed by manually querying the SILVA rRNA database (Quast *et al,* 2013) and extracting the representative sequence from each of our 32 species. Amplicon sequencing reads were then mapped against this database using MAPseq v1.2.4 (Matias Rodrigues *et al,* 2017). Paired reads were mapped independently and assignments were only considered upon agreement. Abundance estimates were then produced by counting the number of reads mapping to each genome included in the study. For metagenome derived estimates, total counts were normalized by the size of the genome (number of base-pairs). For metatranscriptome derived estimates, additional steps were required. Gene predictions by Prokka/Prodigal were used to calculate the total number of coding bases per genome, after exclusion of rRNA regions. Finally, total read counts were normalized by the number of coding bases on each genome.

Species abundance was estimated from metaproteomic data by summing up all filtered protein intensities detected per each species, and dividing the sum by the total summed protein intensity in a given sample.

In addition, to the approaches based on read mapping, several popular tools were used to estimate species abundance. For amplicon sequencing, DADA2 v1.10 (Callahan *et al,* 2016) was used with the GTDB database release 86 (Parks *et al*, 2018) for sequence classification which was limited to genus level classification. Metagenomic and metatranscriptomic species abundances were estimated using mOTUs v2.5 (Milanese *et al,* 2019) and MetaPhlAn v3 (Beghini *et al*, 2021).

### Coverage analyses

Gene, transcript and protein coverage were defined as the number of genes/transcripts/proteins that showed a count higher than 0, divided by the total number of predicted genes per species. For pathway coverage, the same approach was used, but genes/transcripts/proteins were grouped by KEGG pathways instead and thus divided by the number of KEGG orthologs in one single pathway. The same procedure was repeated for metabolites, but using the number of metabolites per pathway as predicted by KEGG instead of the number of KEGG orthologs.

### Mantel test

Mantel test was performed to compare two different kinds of omics datasets and evaluate the similarity between them. Abundance tables of each omics were transformed into distance matrices using 1 - Spearman’s correlation coefficient, and the matrices were compared using the mantel function in the vegan package (version 2.5.5) with the default option. Sixty-one samples that were common among all the omics datasets were used in this analysis.

### Differential species abundance analysis

Differential analysis of species abundance across conditions was performed with ANCOM v. 2.1. Tables of species abundances calculated from each omics measurements were preprocessed with feature_table_pre_process with sample names used as sample variables, condition used as group variable, and parameters out_cut = 0.05; zero_cut = 0.90; lib_cut = 0; neg_lb = TRUE. The ANCOM function was applied to each preproccessed table with condition used as the main variable and time used as the formula for adjustment. P-values were adjusted with Benjamini-Hochberg method (p_adj_method = “BH”). The cutoff of 0.7 for the W statistic was used to identify significantly differentially abundant species (detected_0.7=TRUE).

### Differential transcript, protein and metabolite abundance analysis

Differential transcript analysis was performed using DESeq2 v1.26.0 (Love *et al,* 2014) after taxon-specific scaling (see above). The design formula included the factors run, drug, time point and the interaction term drug:timepoint. Statistical testing was performed with the Wald-test and IHW (Ignatiadis *et al,* 2016) to control the false discovery rate.

Differential protein and metabolite analysis were performed using repeated measures analysis of variance using the lmer function in the ade4 package. The same formula used in the differential transcript analysis was also used in the analysis. To exclude low abundant features, those that have 0 or NA in at least half of the samples were removed prior to the analysis. P-values were adjusted by the IHW method. Fold changes of proteins and metabolites compared to those of controls were calculated based on raw values.

### Pathway and COG enrichment analysis

Pathway enrichment was performed in on differentially abundant features (cutoff for metatranscriptomics abs(log2(fold change))>2, pFDR<0.001, cutoff for metabolomics and metaproteomics abs(log2(fold change))>log2(1.5), pFDR<0.05) with Fisher exact test using stats.fisher_exact in Python 3.7.7. P-values were adjusted with Benjamini-Hochberg procedure with multipletests function from statsmodels. COG enrichment was performed in the R environment using ClusterProfiler (Wu *et al*, 2021).

## Data and code availability

The MS-based proteomics data have been deposited to the ProteomeXchange Consortium via the PRIDE partner repository and are available via ProteomeXchange with identifier PXD036445. Metabolomic data has been submitted to MetaboLights under accession number MTBLS3129. Sequencing data is deposited at the European Nucleotide Archive (ENA): PRJEB46619. Preproccessed data files and tables are available on Figshare at https://doi.org/10.6084/m9.figshare.21667763, and code to generate all figures is available at https://github.com/grp-bork/multiomics_Wuyts_2022.

## Acknowledgements

We acknowledge Vladimir Benes, Matthew Hayward, Melanie Tramontano, Thea Van Rossum, Camille Goemans, Carlos Voogdt and Michael Zimmermann for helpful discussions.

## Funding

The work was supported by the European Molecular Biology Laboratory and has received funding from the European Union’s Horizon 2020 research and innovation programme under grant agreement number 668031 (to RA, SN, SW). KRP: UK Medical Research Council (project number MC_UU_00025/11). MZ-K: Postdoc Mobility Fellowship from the Swiss National Science Foundation (P400PB_186795) and a postdoctoral fellowship from the AXA Research Fund. LM, SGS and TW were supported by the EMBL Interdisciplinary Postdoc (EIPOD) program under Marie Sklodowska-Curie Actions COFUND (grant numbers 291772 and 664726).

## Author Contributions

Experiment design: RA, MD, EK, LM, ATy, KRP, MK, PB. Data analysis: SW, RA, MZ-K, SN, TSBS, MK. *In vitro* pilot studies: MD, EK, TW. *In vitro* experiments: SB, EK, SGS. Metabolomics measurements: DCS. Metagenomic and metatranscriptomic sequencing: EK, RH, ATe. Metaproteomics measurements: PEG, JBM, PVT, MM. Manuscript writing and editing: SW, RA, MZ-K, SN, ATy, KRP, MK, PB.

## Notes

### Competing Interest Statement

The authors have declared no competing interest.

