## Supplementary Figures for "Consistency across multi-omics layers in a drug-perturbed gut microbial community"

### Supplementary Figure 1

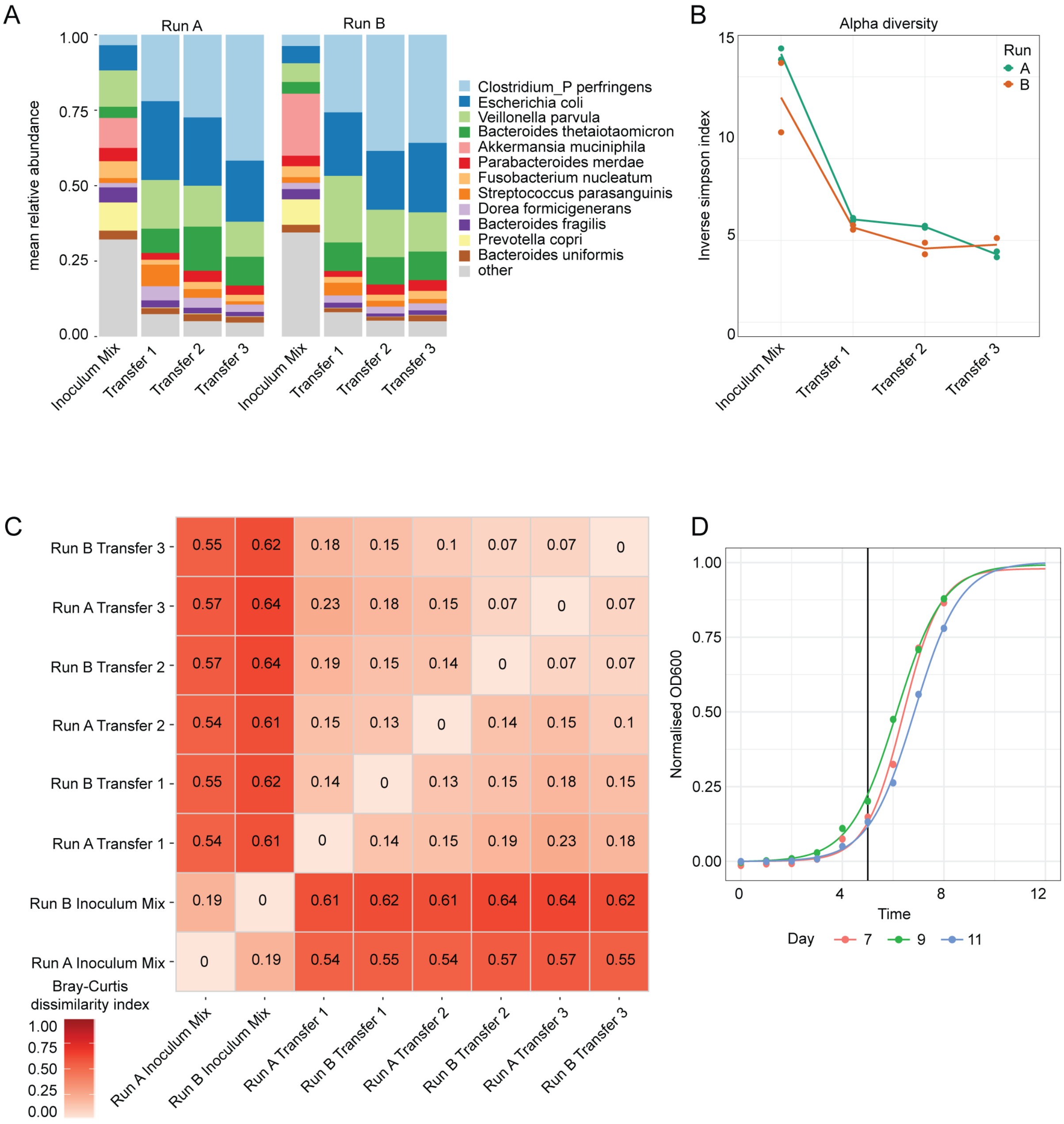


**Supplementary Figure 1. Establishment of a stable microbial community after three community transfers.**

A. Relative abundance of the community members during the community transfer phase prior to drug treatment.

B. Alpha-diversity measurements during the community transfer phase prior to drug treatment.

C. Bray-Curtis dissimilarity values for pairwise comparison of community compositions during the community transfer phase prior to drug treatment.

D. Growth curves were measured every hour during community establishment. We fit a sigmoid function to the measurements per day, and normalised the resulting OD curves. Based on the observed growth curve, we chose to treat the community after five hours (black vertical line) so that the tightly spaced time points within three hours are all within the exponential phase.

**Supplementary Figure 2.**

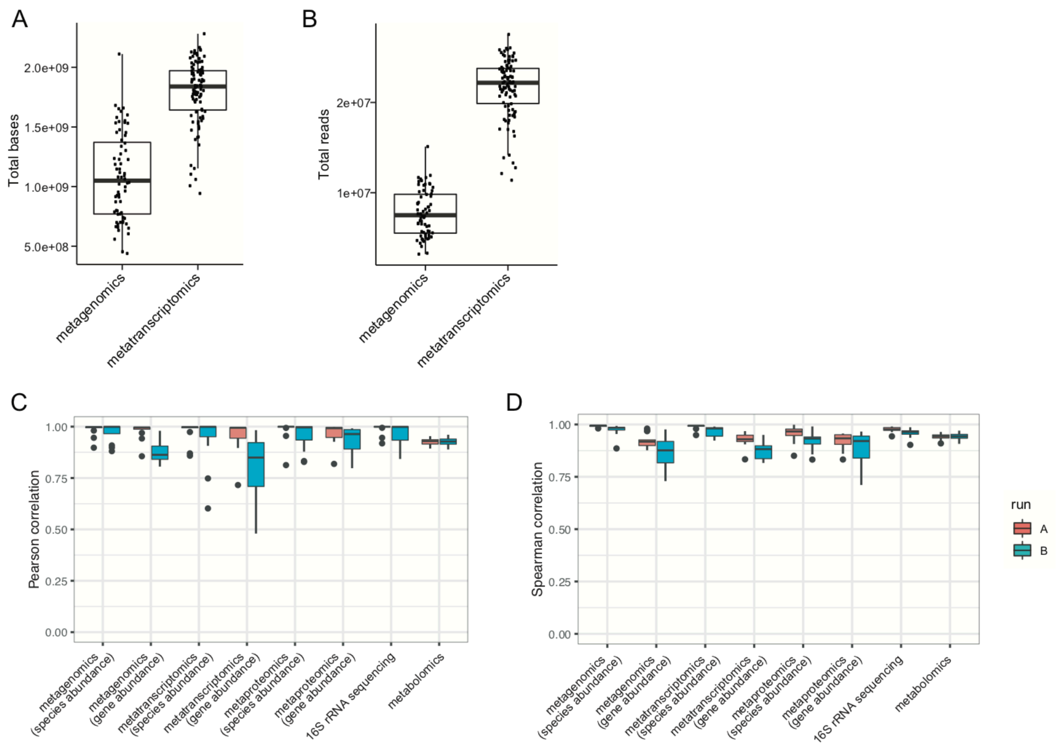


**Supplementary Figure 2. High technical reproducibility across all omics methods.**

A. Total bases detected across metagenomics and metatranscriptomic samples.
B. Total reads detected across metagenomics and metatranscriptomic samples.
C. Pearson correlation coefficients between technical replicates in runs A and B for each omics measurement.
D. Spearman correlation coefficients between technical replicates in runs A and B for each omics measurement.

### Supplementary Figure 3

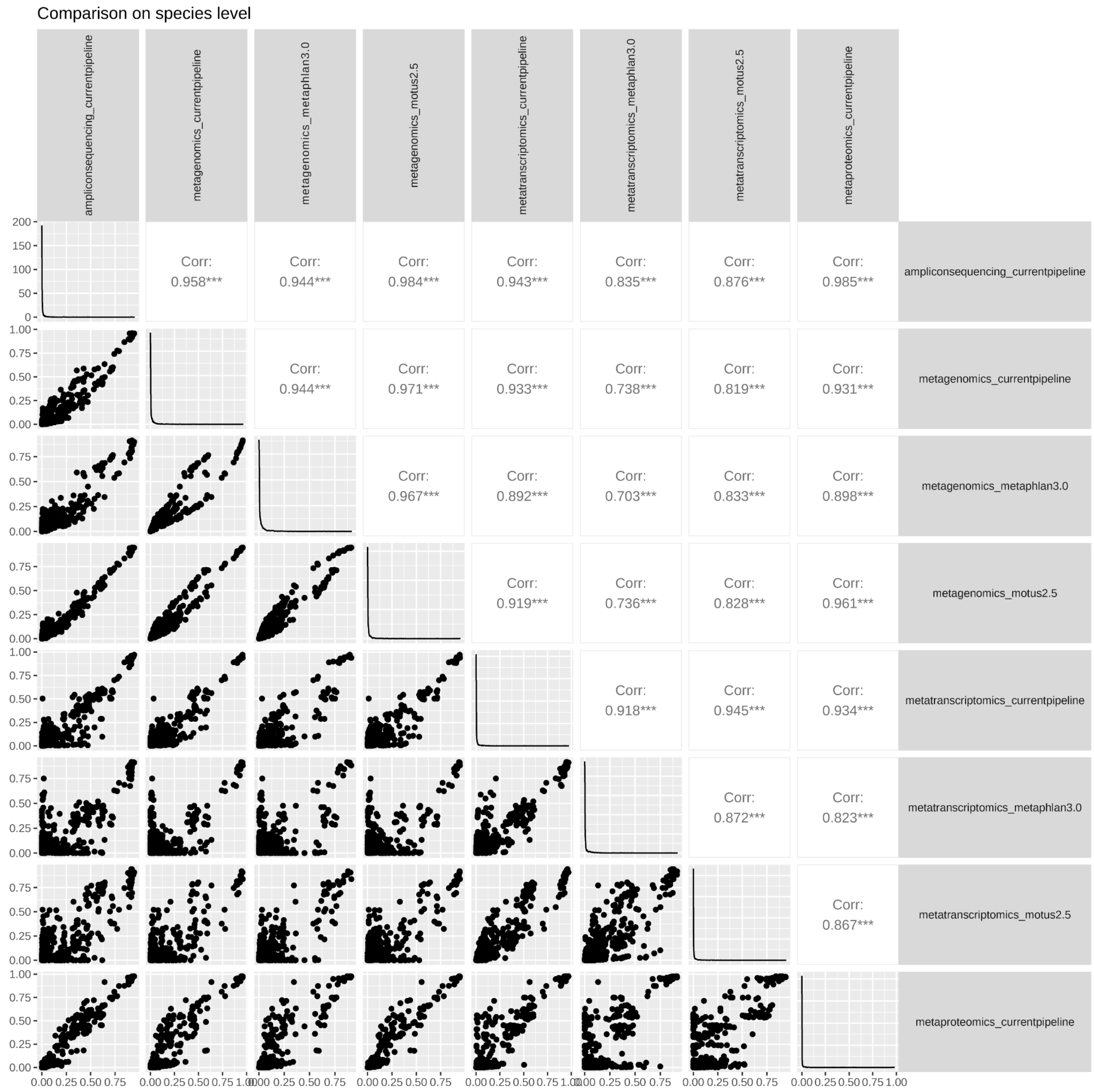


**Supplementary Figure 3. Comparison of species abundance estimation by different pipelines.**

Each dot depicts abundance of one species in one of the four conditions and time points estimated by different methods.

### Supplementary Figure 4

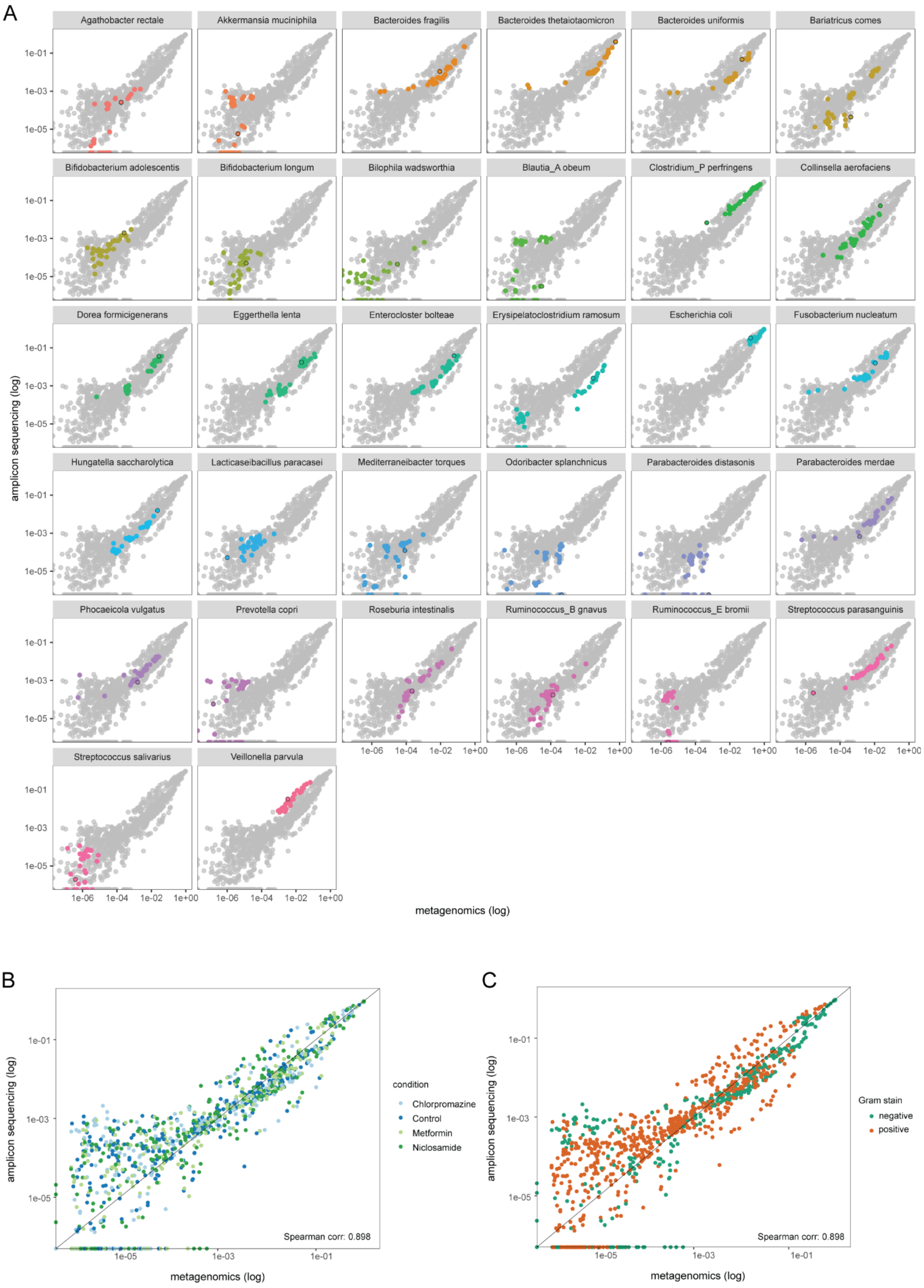

**Supplementary Figure 4. Differences between species abundances estimated by metagenomics and 16S sequencing are not species-, condition-, or gram-type specific.**

A. Metagenomics vs 16S sequencing species abundances coloured by species.

B. Metagenomics vs 16S sequencing species abundances coloured by condition.

C. Metagenomics vs 16S sequencing species abundances coloured by Gram staining.

**Supplementary Figure 5.**

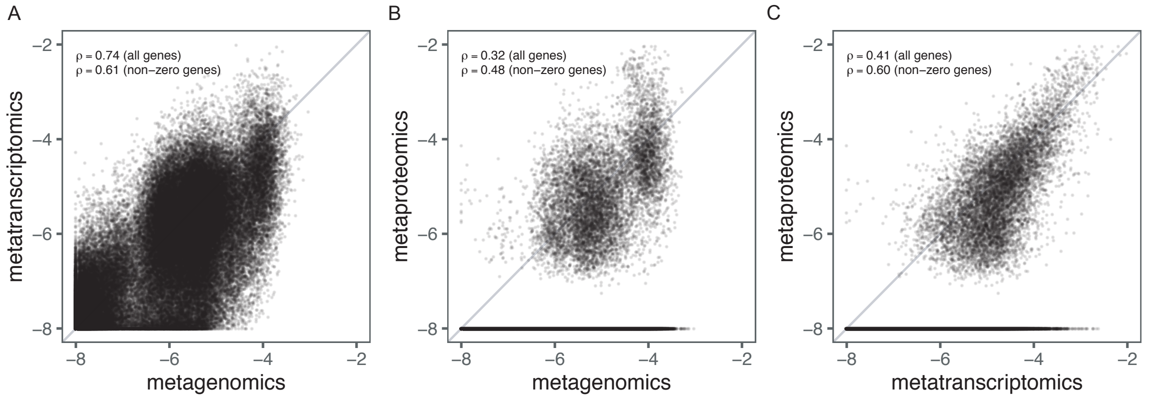


**Supplementary Figure 5. Comparison of gene abundance estimates between different omics.**

Each point corresponds to one gene estimate in one of the four conditions, as detected by A. metagenomics vs metatranscriptomics, B. metagenomics vs metaproteomics, or C. metatranscriptomics vs metaproteomics measurements.

### Supplementary Figure 6

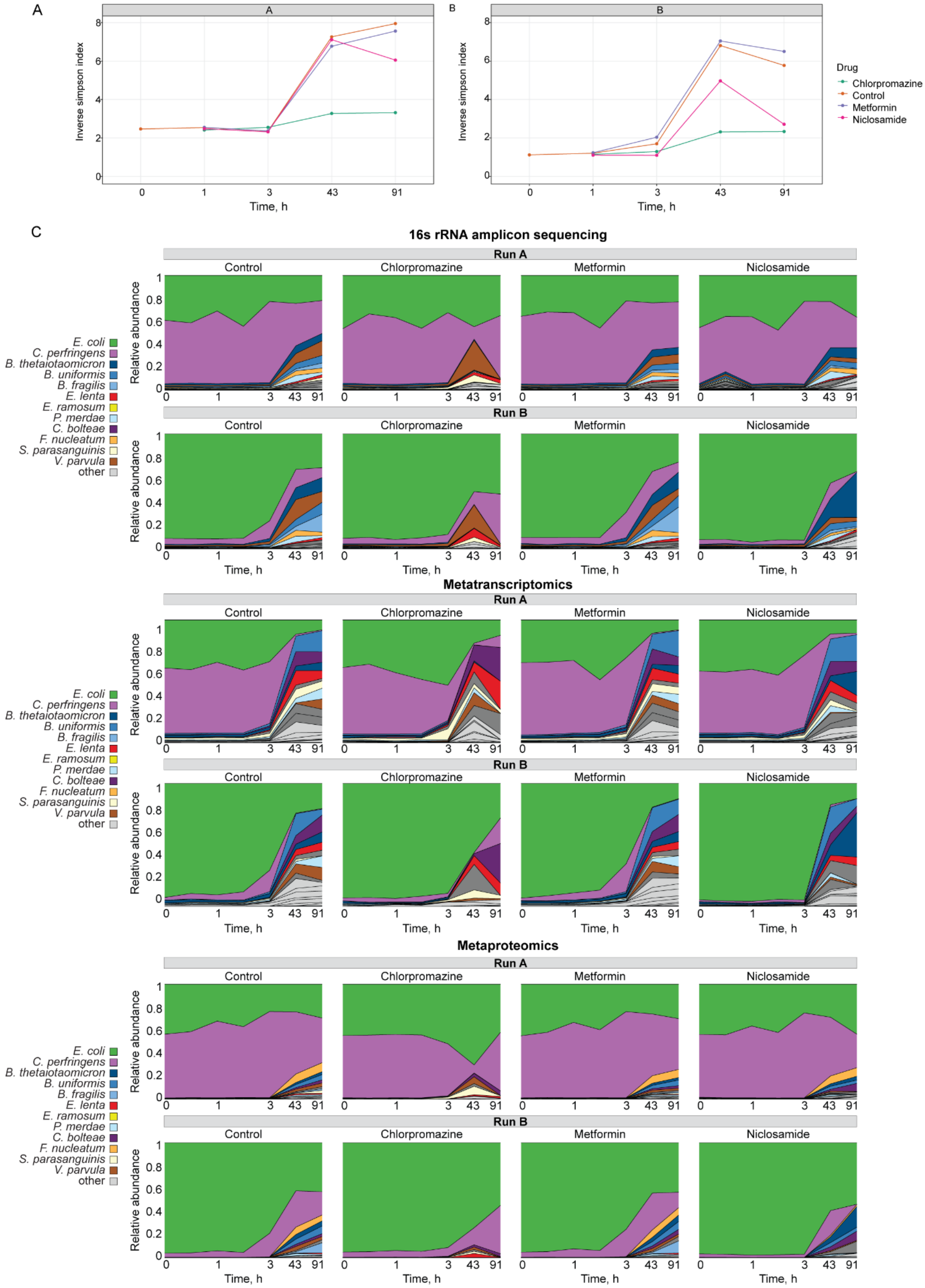


**Supplementary Figure 6. Chlorpromazine strongly affects community composition.**

A., B. Community alpha diversity measurements over time after drug treatment for runs A and B, correspondingly.

C. Relative species abundance changes over time in the three drug conditions and control. Relative abundance measured from 16S rRNA amplicon sequencing, metatranscriptomic and metaproteomic data.

**Supplementary Figure 7**

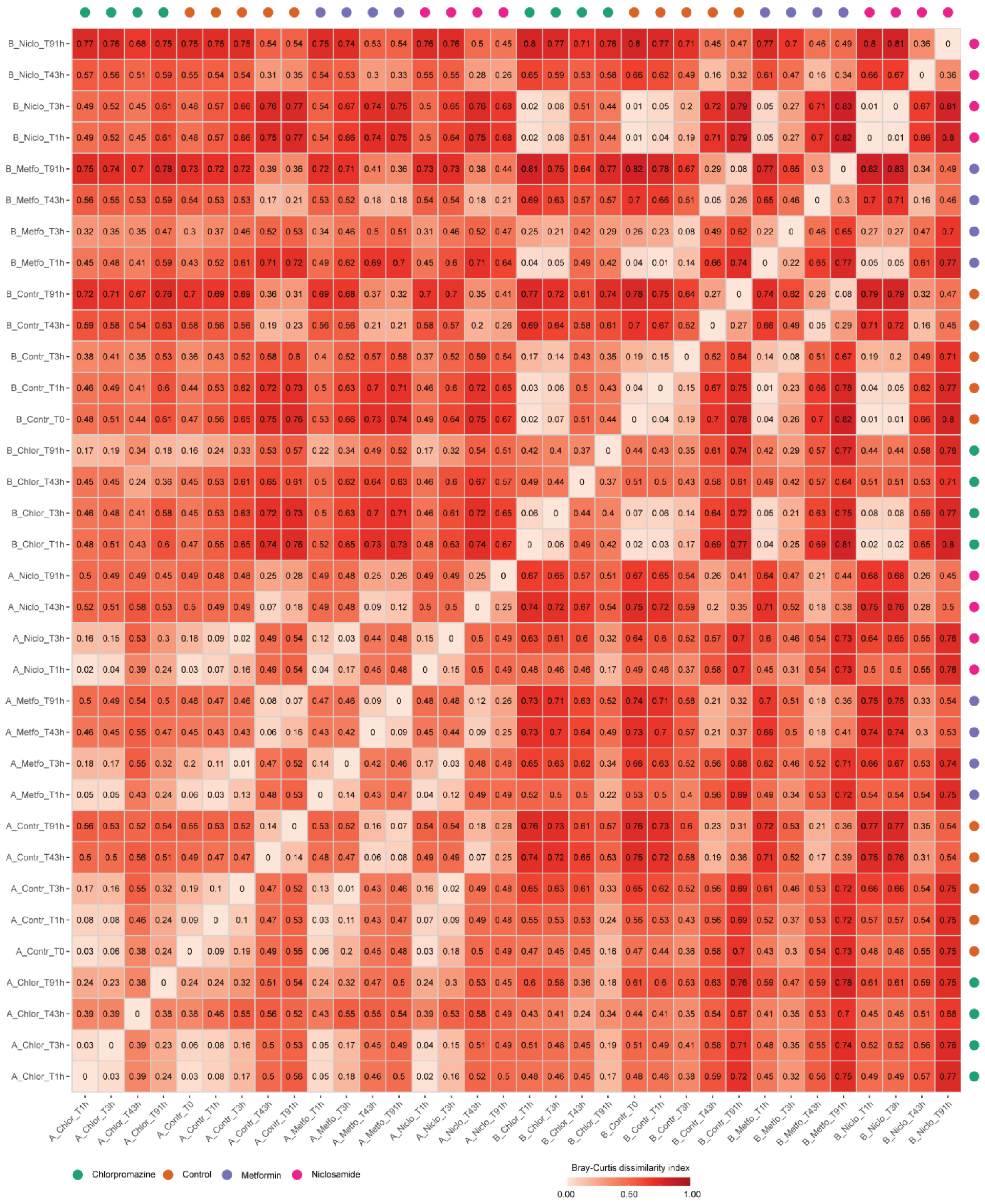

**Supplementary Figure 7. Community composition in different conditions becomes more similar at later time points.**

Bray-Curtis dissimilarity values for pairwise comparison of community compositions at each time point after each of the drug treatments in runs A and B.

**Supplementary Figure 8**

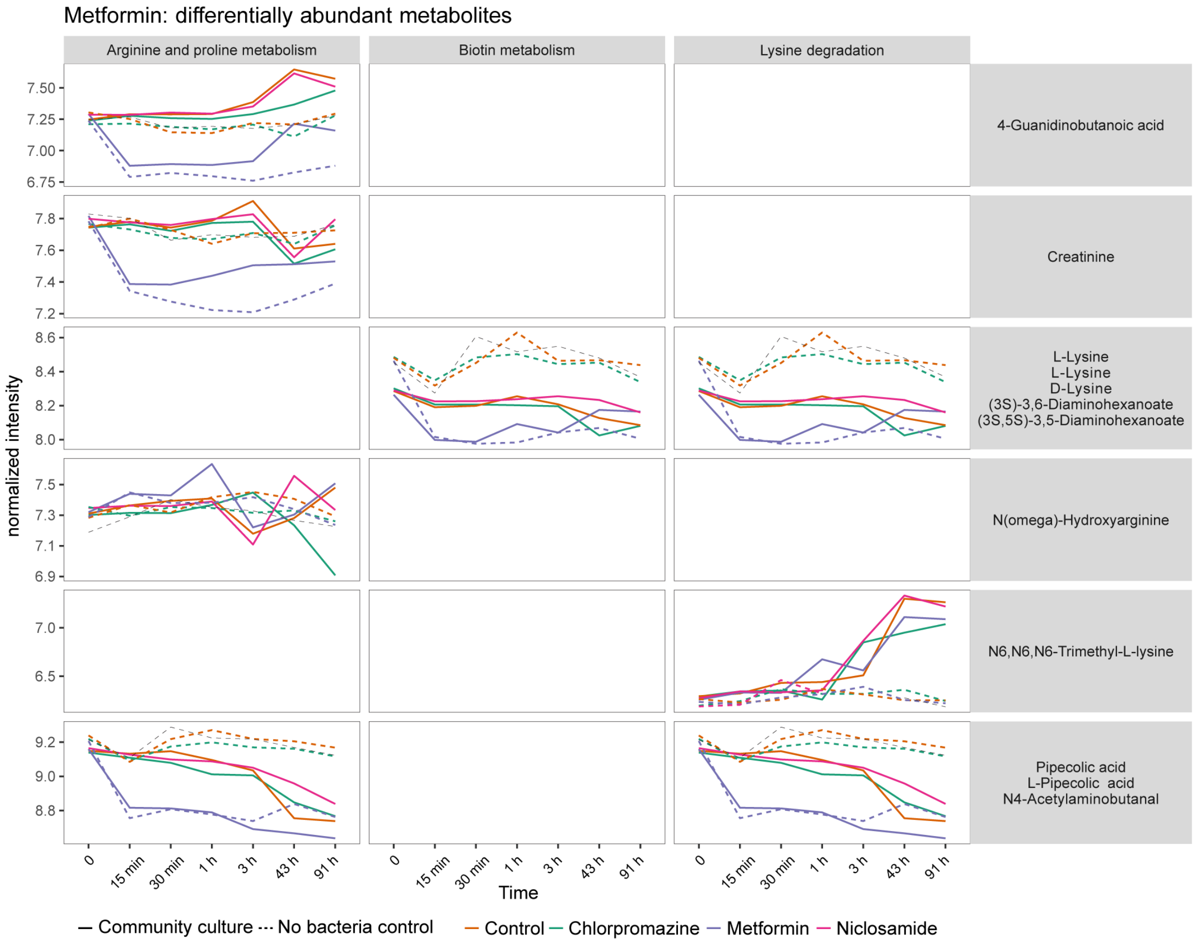


**Supplementary Figure 8. Metabolites increased upon metformin exposure are likely to be measurement artifacts.**

Metabolite profiles overtime after drug treatment in community samples and non-bacteria controls. Metabolites depicted were selected from pathways significantly differentially abundant on metformin revealed by pathway enrichment analysis.

**Supplementary Figure 9**

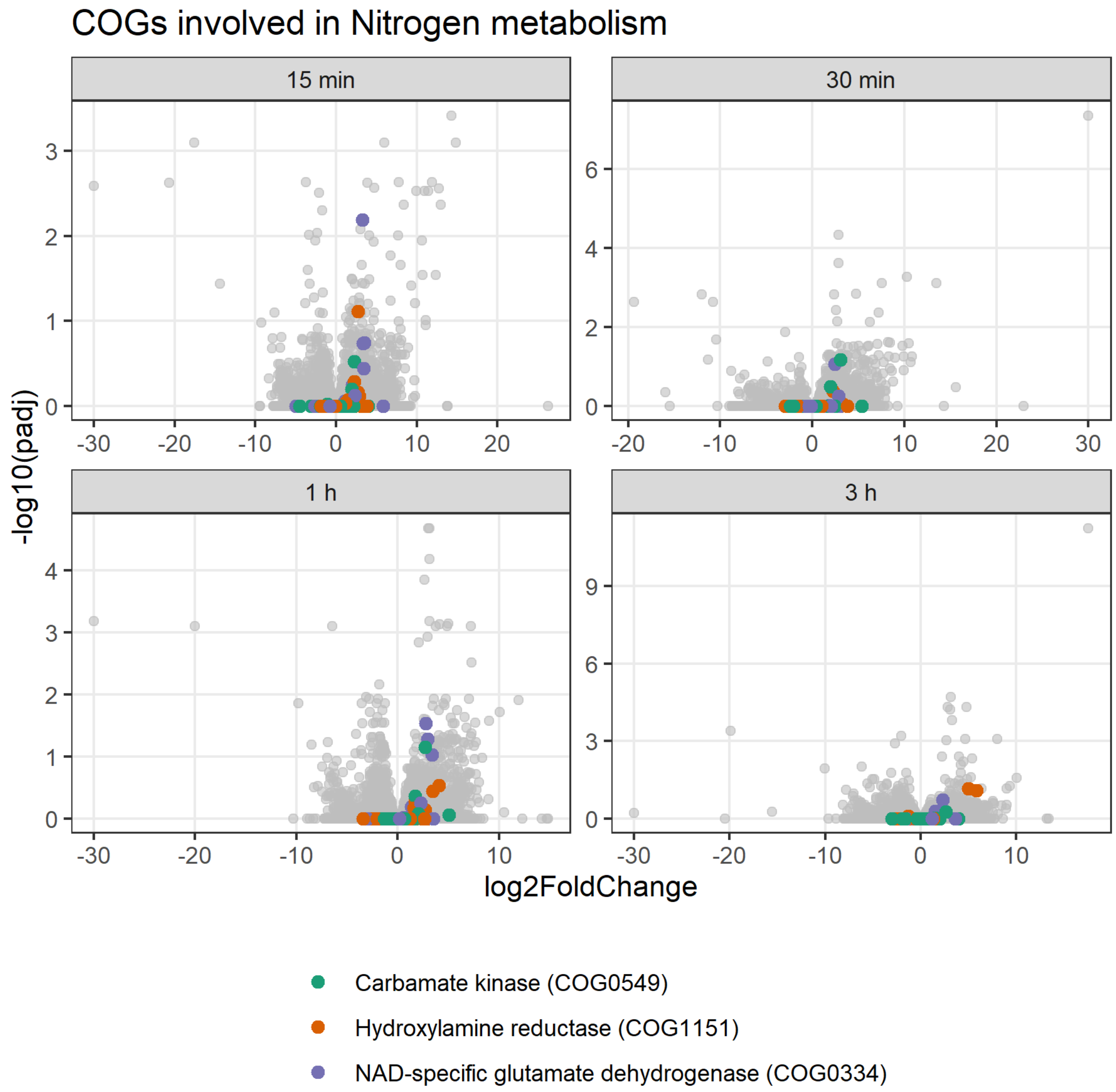


**Supplementary Figure 9. COG categories of genes regulated during early exposure to drugs.**

Volcano plots depicting fold change (log2 scale) and significance (-log10 FDR) of COG abundance upon niclosamide treatment in the first four time points.

**Supplementary Figure 10**

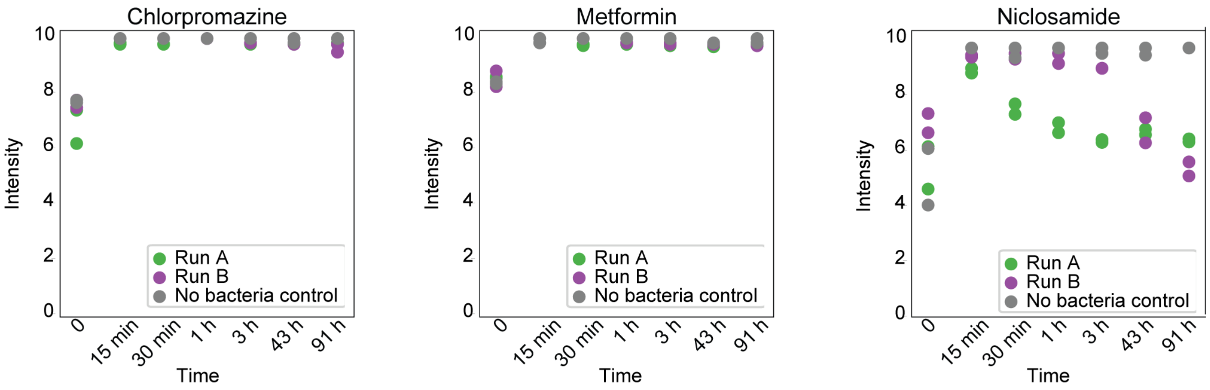


**Supplementary Figure 10. Drug profiles measured over time.**

### Supplementary Figure 11

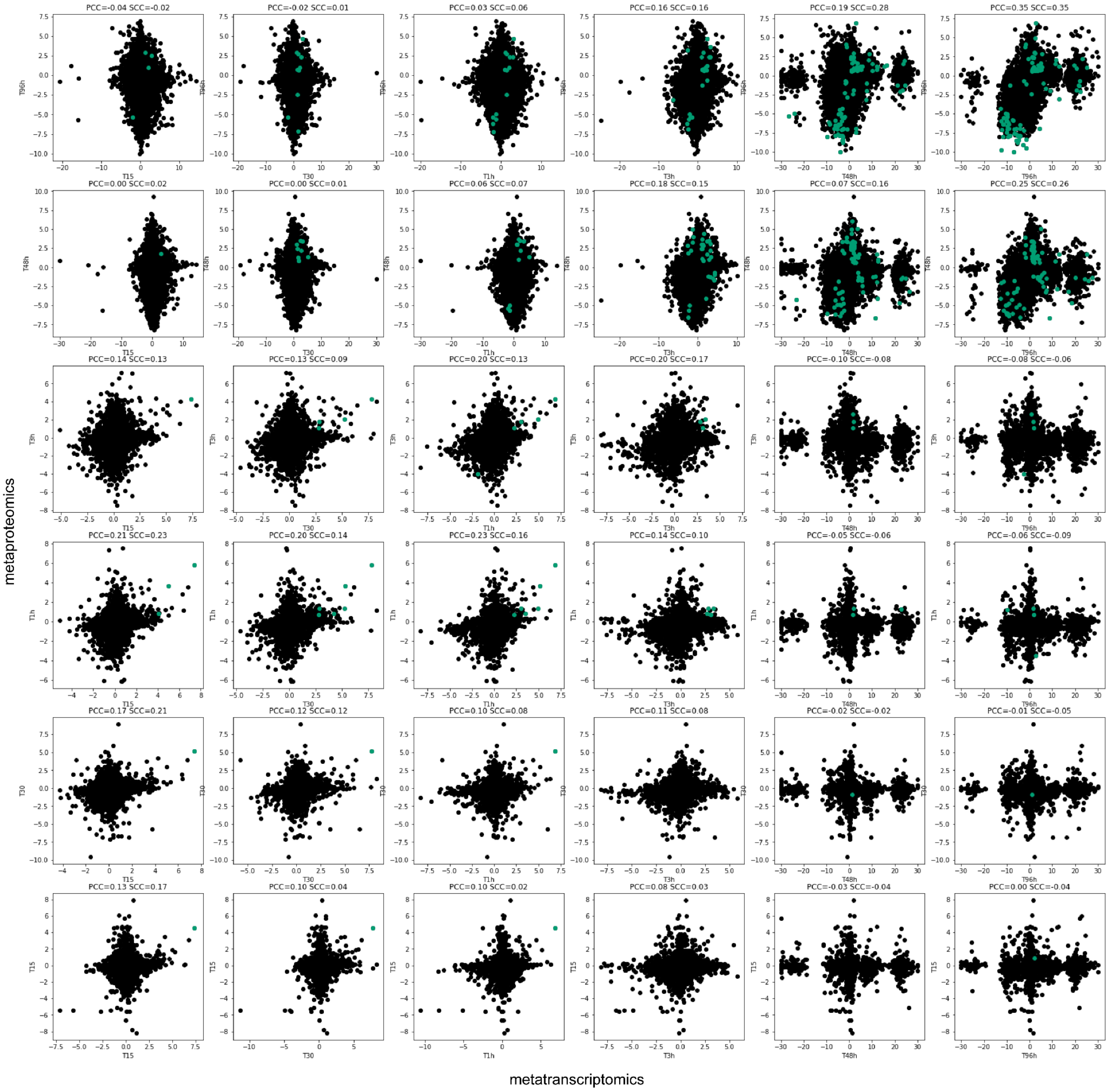


**Supplementary Figure 11. Correlation between transcript and protein fold changes across time points.**

Only proteins detected both by metatranscriptomics and metaproteomics are depicted. Proteins passing the significance FDR cutoff of 0.1 both in proteomics and transcriptomics measurements are depicted in green. PCC - Pearson correlation coefficient, SCC - Spearman correlation coefficient.

**Supplementary Figure 12.**

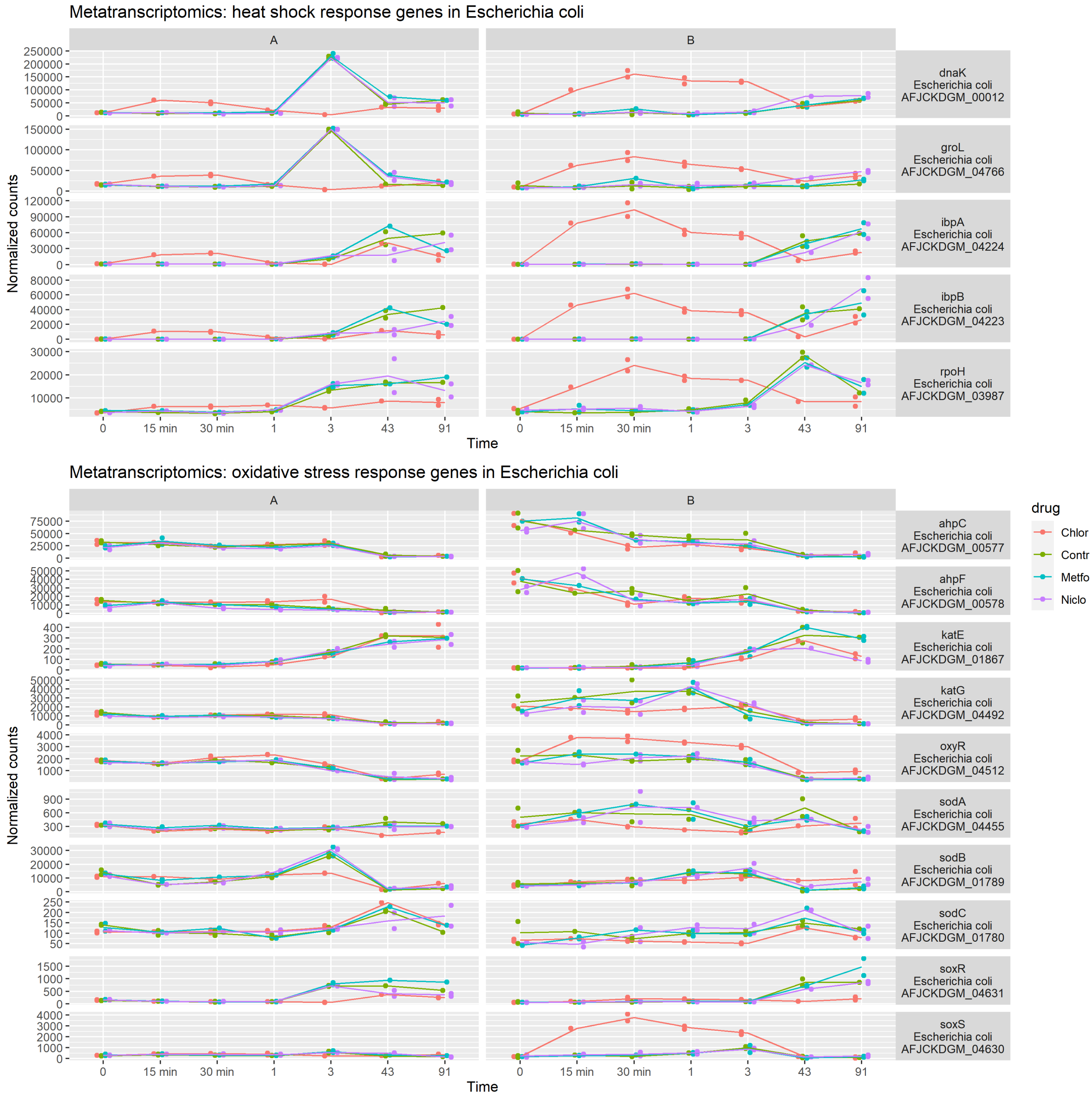


**Supplementary Figure 12. Chlorpromazine causes upregulation of heat shock related genes in *Escherichia coli.* Only a few genes related to oxidative stress are upregulated.**

**Supplementary Figure 13.**

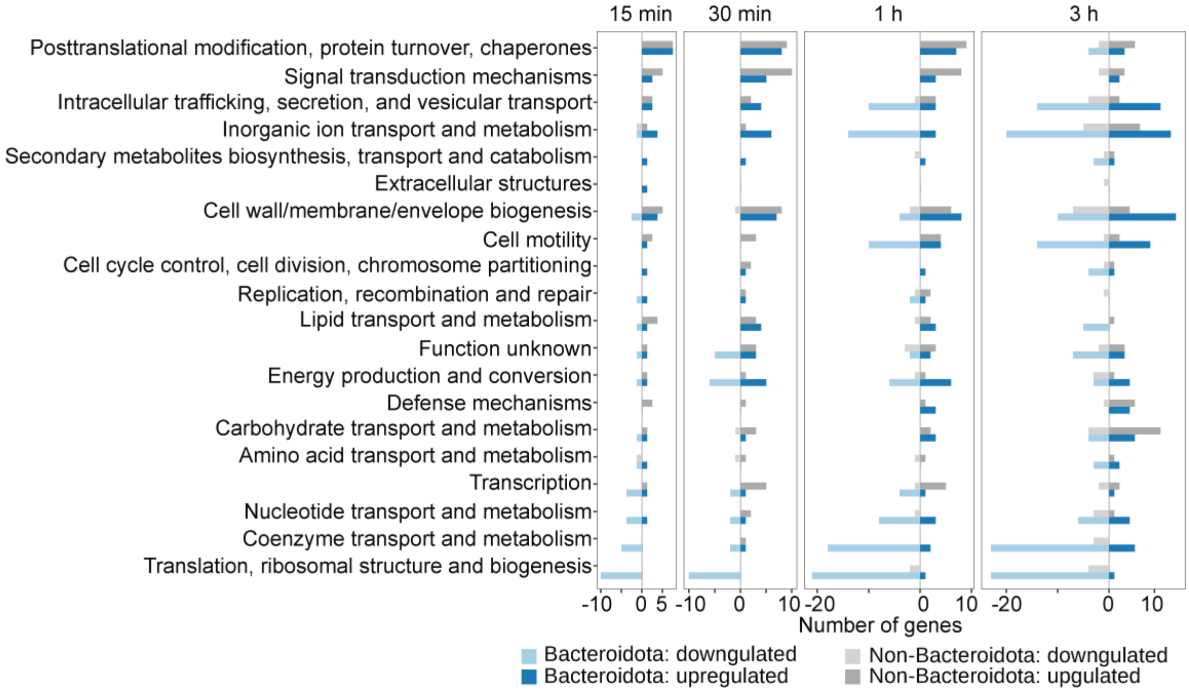


**Supplementary Figure 13. Number of proteins changing in each COG category per time point upon chlorpromazine treatment.**
